# Parsing glomerular and tubular structure variability in high-throughput kidney organoid culture

**DOI:** 10.1101/2025.05.05.652160

**Authors:** Kristiina Uusi-Rauva, Anniina Pirttiniemi, Antti Hassinen, Ras Trokovic, Sanna Lehtonen, Jukka Kallijärvi, Markku Lehto, Vineta Fellman, Per-Henrik Groop

## Abstract

High variability in stem cell research is a well-known limiting phenomenon, with technical variation across experiments and laboratories often surpassing variation caused by genotypic effects of induced pluripotent stem cell (iPSC) lines. Evaluation of kidney organoid protocols and culture conditions across laboratories remains scarce in the literature. We used the original air-medium interface protocol to evaluate kidney organoid success rate and reproducibility with several human iPSC lines, including a novel patient-derived GRACILE syndrome iPSC line. Organoid morphology was assessed with light microscopy and immunofluorescence-stained maturing glomerular and tubular structures. The protocol was further adapted to four microplate-based high-throughput approaches utilizing spheroid culture steps. Quantitative high-content screening analysis of the nephrin-positive podocytes and ECAD-positive tubular cells revealed that the choice of approach and culture conditions were significantly associated with structure development. The culture approach, iPSC line, experimental replication, and initial cell number explained 35-77% of the variability in the logit-transformed proportion of nephrin and ECAD-positive area, when fitted into multiple linear models. Our study highlights the benefits of high-throughput culture and multivariate techniques to better distinguishing sources of technical and biological variation in morphological analysis of organoids. Our microplate-based high-throughput approach is easily adaptable for other laboratories to combat organoid size variability.

## Introduction

Human induced pluripotent stem cell (hiPSC) derived kidney organoids represent an attractive micro-scale source of patient-specific kidney tissues, which are otherwise difficult to obtain^1,2^. Although structurally and functionally immature, PSC-derived kidney organoids have been shown to mimic embryonic kidney development remarkably up to the second trimester, resulting in tubular and glomerular structures with foot processes and a primitive endothelial network with architectural resemblance to nephron structures *in vivo*^3–5^. Initial reports on the differentiation of human pluripotent stem cells (PSCs), which include hiPSCs and human embryonic stem cells (hESCs), into kidney cells and kidney organoids^4,6–9^ were soon followed by studies aiming to enhance the level of maturation and complexity of generated structures, as well as efficiency and cost-effectiveness of the production^10–17^.

However, deployment of a kidney organoid protocol, or any PSC differentiation, can be challenging when the technology is transferred from one laboratory to another. When we set out to establish kidney organoid differentiation in our laboratory and to assess the possibilities to adapt published protocols to higher throughput, it became clear that the differentiation capacity of available cell lines, the choice of protocol, and culture conditions may greatly affect the end-result of the differentiation^18–20^. While it has been reported that laboratory-based variation can surpass genotypic effects, due to issues such as organoid cell type heterogeneity, third-party testing of available methodology, high-throughput applications in particular, remains scarce in the kidney organoid literature^11,20–25^.

The aims of the present study were to evaluate the success and reproducibility of kidney organoid differentiation across available human stem cell lines using a previously published kidney organoid protocol^4^. We further aimed to enhance the high-throughput generation and morphological assessment of kidney organoids, thereby promoting validation, transparency, and transferability of kidney organoid technologies across laboratories.

We set out to test several iPSC lines that were not preselected based on their capacity to differentiate towards kidney cells. The present study largely utilized previously reported healthy donor iPSC lines, but the array of cell lines was complemented by novel iPSC lines generated and characterized in the present study. These include patient-derived iPSC lines of the GRACILE syndrome, a rare early-onset and lethal mitochondrial disorder manifested by fetal growth restriction, aminoaciduria, cholestasis, iron overload in the liver, lactic acidosis, and early death, from which the disease acronym was constructed^26,27^. For kidney organoid differentiation we used one of the first originally described protocols^4,6,9^, a 6-well format protocol by Takasato *et al.,* allowing the production of self-organized organoids generated in albumin polyvinyl alcohol essential lipids (APEL) medium at air-medium interface (AMI) (“APEL/air-medium interface protocol”)^4^ (Figure 1A).

**Figure 1.**
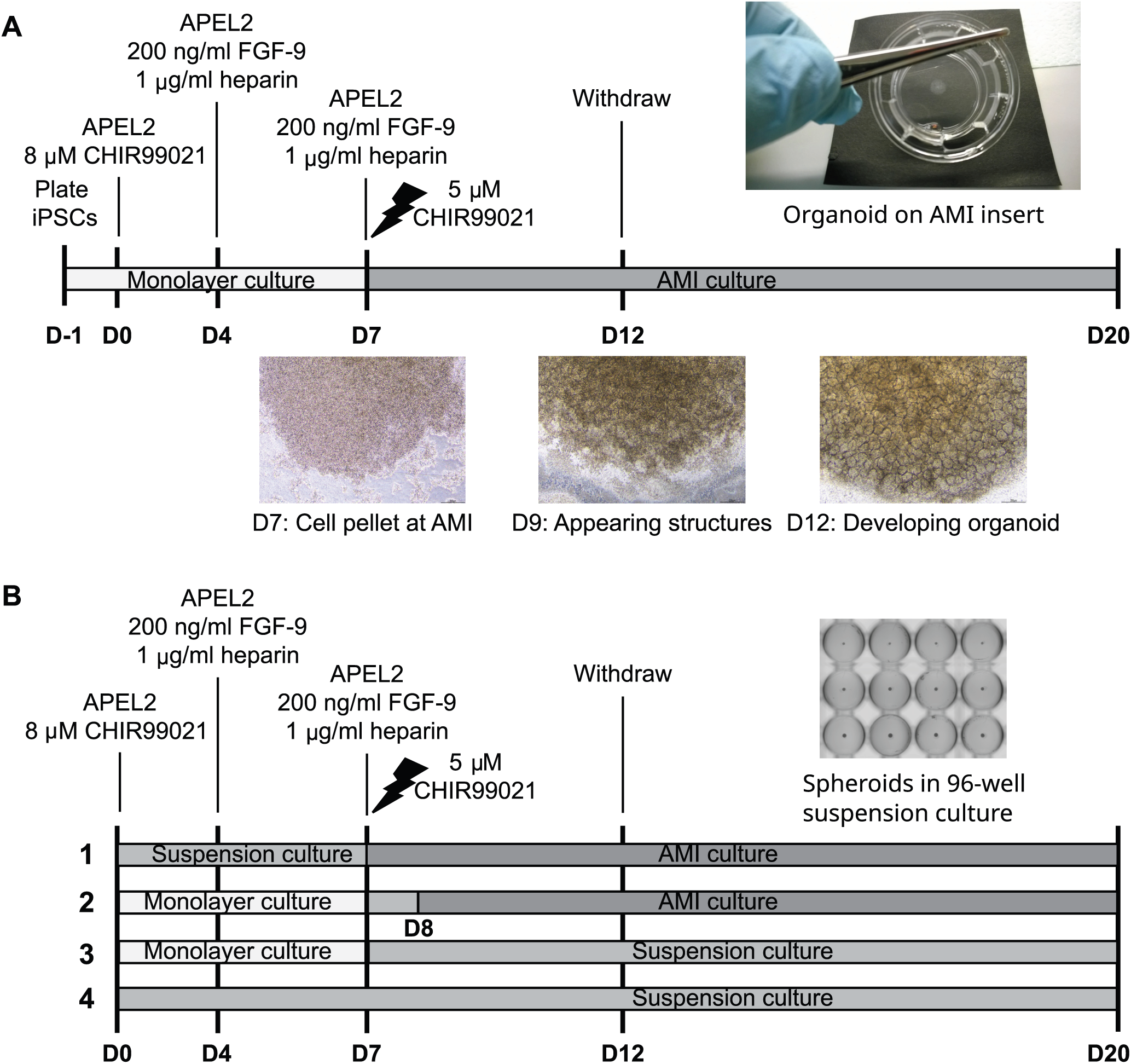
Overview of the protocols utilized in the generation of iPSC-derived kidney organoids. (A) The previously established protocol, a 6-well format air-medium interface (AMI) system utilizing albumin polyvinyl alcohol essential lipids medium 2 (APEL2) as a basal medium throughout the experiment (the “APEL/AMI protocol”) was tested for the success rates and reproducibility of iPSC-derived kidney organoids. (B) The original APEL/AMI protocol was modified towards higher throughput by testing different culture systems, which incorporate spheroid suspension culture steps into the protocol (approaches 1–4). D, day; FGF-9, fibroblasts growth factor 9.

Respective success rates and reproducibility of organoid generation were determined in a descriptive manner based on the appearance of intrinsic structures in light and immunofluorescence microscopy. Subsequently, the original kidney organoid protocol was modified to incorporate spheroid culture steps in four different high-throughput approaches utilizing multiwell plates (Figure 1B). The proportion of immunostained nephrin-positive glomerular structures and epithelial cadherin (ECAD)-positive tubular structures were assessed with a quantitative high-content confocal screening system to set up a more efficient and consistent protocol to fulfil the requirements of high-throughput analyses.

## Materials and methods

### Ethics

The present study was performed as a collaboration between Folkhälsan Research Center and University of Helsinki and conducted in accordance with the Declaration of Helsinki. The study was carried out from October 2014 onward and benefited from previously collected human fibroblasts and reprogrammed iPSC lines. All ethical approval statements were obtained from the ethics committee of the Helsinki and Uusimaa Hospital District, Finland (HUS; 77/E7/2007, obtained in 2007, for GRACILE patient fibroblasts; an update of 77/E7/2007, obtained in 2011, for GRACILE iPSCs and differentiation; 423/13/03/00/08, obtained in 2009, for healthy donor fibroblasts/iPSCs and differentiation). Written informed consent for participation were obtained from the donor or the guardians. Healthy donor fibroblasts were collected, and respective iPSC reprogramming experiments were started between January 2010 and May 2012. GRACILE samples were collected, and respective iPSC reprogramming experiments were started between April 2007 and September 2011. The authors had no access to information that could identify individual participants during the study.

### Cell culture

Human iPSC lines HEL11.4, HEL24.3, and HEL47.2 from healthy donors have been generated and described previously^28–30^. hESC line H9 (WA09) was originally from WiCell and Dr. James Thomson, University of Wisconsin and provided by BSCC, Helsinki, Finland. Healthy control iPSC line HEL61 and GRACILE syndrome HEL124 were generated in this study.

PSCs were cultured on Matrigel-coated (BD Biosciences) cell culture plates in E8 medium (Life Technologies) and split using 0.5 mM ethylenediaminetetraacetic acid (EDTA; Thermo Fisher Scientific). Rho-associated protein kinase (ROCK) inhibitor (Y-27632; Sigma) was utilized to enhance the cell survival after dissociation and thawing (5.0–10.0 μM). Media were changed every other day. Antibiotics were utilized at routine cell culturing. GRACILE iPSC lines HEL124.1 and HEL124.2 were supplemented with 200 μM uridine. Uridine was also added to the media of all cell cultures in the experiments, where the growth or characteristics of HEL124.1 and HEL124.2 were compared to the healthy donor cell lines. All cells were kept in an incubator at +37°C and 5% CO2. Cell cultures were regularly tested for the mycoplasma infection (LookOut® Mycoplasma qPCR Detection Kit, Sigma).

### Generation and characterization of healthy control iPSC line HEL61 and GRACILE syndrome iPSC line HEL124

iPSC reprogramming was done at Biomedicum Stem Cell Center, University of Helsinki (BSCC; Helsinki, Finland). Dermal fibroblast cultures of a 57-year-old healthy female and a newborn female GRACILE patient (ZCC11-72) were cultured in Dulbecco’s modified Eagle’s medium (DMEM; Life Technologies/Thermo Fisher Scientific) supplemented with 10% fetal bovine serum (FBS; Life Technologies) and 2 mM GlutaMAX (Life Technologies/Thermo Fisher Scientific) (the “fibroblast medium”). The medium of GRACILE fibroblasts was additionally supplemented with 200 μM uridine (Calbiochem).

The fibroblasts from the healthy female were seeded four days before reprogramming. On the day of reprogramming (day 0), cells were dissociated into single cells with trypsin and washed with phosphate-buffered saline (PBS). Cells were transfected using a Sendai virus (SeV) kit according to the manufacturer’s instructions (CytoTune-iPS Reprogramming Kit, Life Technologies) and plated on mitomycin C-inactivated mouse embryonic fibroblasts. After five days, cell culture medium was changed to human embryonic stem cell medium (hESC medium; KnockOut DMEM supplemented with 20% KnockOut-Serum Replacement (KOSR), 1% GlutaMAX, 0.1 mM β-mercaptoethanol, 1% nonessential amino acids; all from Life Technologies) containing 6 ng/ml recombinant human basic fibroblast growth factor (FGF-2; Sigma) and supplemented with 0.25 mM sodium butyrate (Sigma). Emerging colonies were picked manually, plated on Matrigel-coated cell culture plates in Essential-8 medium (E8 medium; Life Technologies), and further expanded for the generation of HEL61 iPSC lines. Media were changed every other day. The clone HEL61.2 was included in the analyses of the study.

The fibroblasts (1x10^5^ cells) of the GRACILE patient were transduced with SeVdp (a single vector harboring the four reprogramming factors) as previously described^31,32^, and plated on Matrigel (BD Biosciences) in the presence of the fibroblast medium supplemented with 200 μM uridine. After five days, cell culture medium was changed to the hESC medium supplemented with 0.25 mM sodium butyrate (Sigma) and 200 μM uridine. When first colonies started to emerge, cell culture medium was changed to E8 medium. iPSC colonies were picked manually and further expanded in the presence of 200 μM uridine. The clones HEL124.1 and HEL124.2 were characterized and the presence of the disease-causing *BCS1L* mutation was verified by Sanger sequencing.

Chromosomal integrity of HEL124.1 and HEL124.2 was analyzed from extracted DNA at the Finnish Microarray and Sequencing Centre (FMSC, Turku, Finland) by a KaryoLite™ BACs-on-Beads™ method (KaryoLite™ BoBs™, Perkin Elmer)^33^ with two technical replicates per sample. The karyotype of HEL61.2 was analyzed by G-banding.

In the characterization of selected iPSC clones, the expression of embryonic stem cell marker proteins TRA-1-60, OCT3/4, and SSEA3 or SSEA4 was assessed by immunofluorescence microscopy of iPSCs, and the expression of embryonic stem cell marker genes *OCT3/4*, *SOX2*, *NANOG*, and *TDGF1* as well as the clearance of the SeV were verified by quantitative reverse transcription PCR (RT-qPCR). Spontaneous differentiation capacity of here generated iPSC lines was tested by embryoid body formation and subsequent immunofluorescence staining of the markers of the three germ layers (α-fetoprotein, β-III tubulin, MAP-2, vimentin, and desmin).

### Reverse transcription and quantitative PCR (qPCR)

Total RNA was extracted using RNA Spin II kit (Macherey-Nagel) according to the manufacturer’s instructions. cDNA was synthesized from 1 μg total RNA by SuperScript III reverse transcriptase (Invitrogen) with oligo dT primer (Invitrogen) in 20 μl volume. One per cent of above cDNA was used for each qPCR reaction in a 20 μl mixture containing 10 μl of SYBR green-Taq mixed solution (Sigma) and 5 μl of 2 μM-primer mix. PCR reactions were carried out in a Corbette thermal cycler (Qiagen) for 40 cycles and each cycle contained +95°C for 15 s, +60°C for 30 s and +72°C for 30 s. Relative gene expression was determined using ΔΔCt method, with a glyceraldehyde-3-phosphate dehydrogenase (*GAPDH*) utilized as a housekeeping gene. Expression levels are relative to the RNA sample of hESC line H9 (positive control). Primer sequences (5′→3′) were as follows: GGATCACTAGGTGATATCGAGC (SeV-F), ACCAGACAAGAGTTTAAGAGATATGTATC (SeV-R); TTGGGCTCGAGAAGGATGTG (OCT4-F), TCCTCTCGTTGTGCATAGTCG (OCT4-R), GCCCTGCAGTACAACTCCAT (SOX2-F), TGCCCTGCTGCGAGTAGGA (SOX2-R), CTCAGCCTCCAGCAGATGC (NANOG-F), TAGATTTCATTCTCTGGTTCTGG (NANOG-R), TCAGAGATGACAGCATTTGGC (TDGF1-F), TTCAGGCAGCAGGTTCTGTTTA (TDGF1-R), GGTCATCCATGACAACTTTGG (GAPDH-F), TGAGCTTCCCGTTCAGCTC (GAPDH-R).

### Embryoid bodies

Cells were dissociated into small aggregates and passaged on ultra-low-attachment 6-well culture plates (Corning) in hESC medium without FGF-2. The embryoid body culture medium was supplemented overnight with 5 µM ROCK inhibitor (Y-27632, Selleckchem or Sigma) after the initial cell plating to improve cell viability. 200 μM uridine was added in the medium of HEL124.1 and HEL124.2 GRACILE cell line cultures. Embryoid bodies were grown in suspension for 4–19 days, and the medium was changed regularly. Thereafter, the embryoid bodies were plated on gelatine-coated coverslips and cultured for additional 6–10 days before processing for immunocytochemistry.

### Immunocytochemistry of iPSCs and embryoid bodies

Cultures were fixed at room temperature with 4% paraformaldehyde (PFA) and permeabilized using 0.2% Triton X-100 in PBS. Non-specific proteins were blocked by Ultra V block (Thermo Fisher Scientific) or 0.5% BSA in PBS. The cells were then incubated with diluted primary antibodies overnight at +4°C or 1 hour at RT. Secondary antibody incubations were done at RT for 40 min in the presence of Hoechst nuclei stain. Antibodies are described in Supplementary Table S1.

### Differentiation of iPSCs towards hepatocyte lineage cells

The protocol utilized for the definitive endodermal differentiation was based on previously published protocol^34^. After modifying the protocol to exclude sodium butyrate and testing cell line specific requirement for insulin, healthy donor iPSC line HEL24.3 and both available GRACILE iPSC clones (HEL124.1 and HEL124.2 representing the same patient, ZCC11-72) were successfully differentiated to definitive endoderm. Differentiation of HEL61.2 required insulin supplementation, which was observed to result in a more heterogenous culture in this particular cell line as compared with the differentiated cultures of other utilized cell lines.

### Western blotting

Samples were lysed in cold 50 mM Tris-HCL pH 7.4, 150 mM NaCl, 1% Triton X-100, 1 mM ethylene glycol tetra acetic acid (EGTA) lysis buffer supplemented with protease inhibitor cocktail (Roche). The cells were disrupted on ice with a pestle, and the final lysate was collected after centrifugation at +4°C for 10 minutes (13 000 rpm in a microcentrifuge). Protein concentrations of the lysates were analyzed by DC Protein Assay (Bio-Rad). Protein lysates (1 or 10 μg) were loaded into single SDS-PAGE gel per comparison (Mini-Protean TGX Any kDA, Bio-Rad), electrophoresed, and semi-dry blotted on 0.2 μm PVDF membrane (Trans Blot Turbo, Bio-Rad). Blots were cut horizontally for simultaneous probing of different proteins (>50 kDa, 37 – 50 kDa, and < 37 kDa). The stainings were carried out according to standard procedure with well-established primary and secondary antibodies (Supplementary Table S2). Stained blots were exposed to HRP substrate (Pierce ECL Western Blotting Substrate or SuperSignal West Femto Maximum Sensitivity Substrate, Thermo Fisher Scientific) and imaged fresh for chemiluminescence (ChemiDoc Imager, Bio-Rad). Chemiluminescence signals were quantified (ChemiDoc Image Lab Software, Bio-Rad) and normalized against the signals of mitochondrial (porin or SDHA) or cytoplasmic (β-actin) loading controls. The relative expressions of samples representing the same genotype (i.e., control or mutated cell lines) were pooled for visualization of the mean results.

### Differentiation of PSCs into kidney organoids

Kidney organoids were generated as described^4,14^ with the following technical modifications and additional notes (Figure 1A). In APEL/air-medium interface differentiation^4,35^, cells were detached with 0.5 mM EDTA (Thermo Fisher Scientific), and small cell aggregates were plated at different densities on Matrigel-coated cell culture dishes in E8 medium (Thermo Fisher Scientific) supplemented with 2.5–5.0 μM ROCK inhibitor (Y-27632, Sigma). Next day, cells were washed, and the differentiation was continued according to the original protocol except that 5% Protein-Free Hybridoma Medium (PFHM-II; Thermo Fisher Scientific) was separately added to the basal medium available at the time of the study (a recombinant protein-based, animal product-free APEL medium, i.e., APEL2; Stem Cell Technologies). Of note, none of the tested cell lines were able to differentiate without PFHM. At day 4, glycogen synthase kinase 3 inhibitor (CHIR99021; Tocris Bioscience) containing medium was changed to animal-free FGF9 (PeproTech) and heparin (Sigma) containing medium according to the original protocol. Approximately 6 × 10^6^ cells were obtained from 12-well (3.8 cm^2^) by day 7 of the differentiation. At day 7, cells were detached and aliquoted in microtubes, in which the one-hour shock with 5 μM CHIR99021 (Tocris Bioscience) in APEL2 medium in a cell incubator was carried out, before the cells were pelleted and placed on Transwell membranes (Corning/Sigma).

The following Transwell membranes were utilized: 10 µm thick polyester membrane with 0.4 µm pores and 4×10^6^/cm^2^ pore density (#3450), and 10 µm thick polycarbonate membrane with 0.4 µm pores and 1×10^8^/cm^2^ pore density (#3412). Kidney organoids were harvested at day 20 unless otherwise stated. Throughout the differentiation, the medium was changed every second day, and the differentiating cells were monitored daily under the light microscope. Uridine was also added to media of all cell cultures in the experiments where the growth or characteristics of the GRACILE cell lines were compared to the healthy donor cell lines.

The following changes allowing higher throughput by incorporating spheroid culture steps, were applied to the APEL/air-medium interface protocol (Figure 1B, approaches 1-4). For spheroid formation, iPSCs were detached with TrypleSelect (Thermo Fisher Scientific), counted, and diluted in different densities (ranging from 2 000 to 50 000 cells for approaches 1 and 4 or 100 000 to 200 000 cells for approaches 2 and 3) in 100 µl of medium per 96-well of a CellCarrier Spheroid Ultra-low attachment (ULA) microplate (PerkinElmer) followed by spinning to enhance cell aggregation/spheroid formation. For the spheroid cultures, the CHIR99021 shock at day 7 was carried out in the ULA wells (Figure 1B, approaches 1 and 4), while the shock for the cells detached from the monolayer cultures, for subsequent spheroid formation, was carried out in a microtube containing a master dilution to be aliquoted in each 96-well (Figure 1B, approaches 2 and 3). For the Opera Phenix high-content screening of structures, approaches 1-4 were tested with two selected healthy-donor iPSC lines, HEL24.3 and HEL61.2, both in two separate experiments (experiment A and B) and 24-well AMI inserts were used in approaches 1 and 2 (Supplementary Table S3).

In the differentiation of approach 1 at AMI on 96-well membranes, an array of 96 wells with permeable inserts connected by a rigid, robotics-friendly tray was utilized [HTS Transwell-96 Permeable Support with 10 µm thick polyester membrane containing 1.0 µm pores (#3380)]. Medium was changed regularly according to the original protocol ^4,35^, except that the medium for the spheroid cultures from day 9 onwards was changed daily.

The same or subsequent batches of each iPSC line were utilized throughout the study, unless otherwise stated. Further information with schematic representations of all kidney organoid protocols utilized in the present study are provided in the results section.

### Immunofluorescence staining of kidney organoids

Samples were fixed with 4 % PFA in PBS supplemented with 1 mM CaCl_2_ and 0.5 mM MgCl_2_ and washed with PBS. Samples differentiated at AMI in a 6-well Transwell inserts (Corning) were detached from the membrane and cut in half for separate staining. The stainings were processed in multiwell plates, except in case of large organoids, which were incubated in small droplets of antibody dilution placed on a clean sealing film. Cells were permeabilized and blocked for unspecific binding with 0.3% Triton X-100 in 10% FBS-PBS, for 4–5 hours at RT. After washing with 0.1% Triton X-100 and 10% FBS in PBS the samples were incubated in primary antibody dilution, at +4°C overnight. Fluorescein Lotus tetragonolobus lectin (LTL; Vector Laboratories FL-1321, 1:500) and following primary antibodies were utilized: guinea pig anti-mouse nephrin (Progen GP-N2, 1:300) and mouse anti-human epithelial cadherin (ECAD; BD Biosciences 610181, 1:400). After washing, the samples were incubated in a secondary antibody dilution in dark, for 4–5 hours at RT or alternatively, at +4°C overnight. The following secondary antibodies were utilized: Goat anti-Guinea pig AlexaFluor 488 (Thermo A11073, 1:1 000), and goat anti-mouse AlexaFluor 594 (Thermo A11005, 1:1 000). Finally, samples were washed, and nuclei were stained (Hoechst 33258, Sigma), followed by rinsing with lab-quality water and mounting with in-house-made Mowiol mounting medium or Fluoromount Aqueous Mounting Medium (Sigma).

### Microscopy and morphological analyses of kidney organoids

Whole mount organoids were viewed and imaged for overall appearance using the Nikon Eclipse Ts2 light microscope and DS-Fi3 camera with DS-L4 imaging application. The immunostained organoids were viewed and imaged using an Axioplan 2 microscope and the AxioCam HRc camera with an AxioVision imaging system (Carl Zeiss Microscopy GmbH), and additionally, using Axio Imager M2, AxioCam 503, and ZEN 2.3 software. Kidney organoids were also imaged using PerkinElmer Opera Phenix spinning disk confocal microscope.

### Opera Phenix high-content screening and image analysis

High-content imaging of whole mount organoids was performed with a PerkinElmer Opera Phenix spinning disk confocal microscope (High Content Imaging and Analysis Unit, FIMM, HiLIFE, University of Helsinki, Finland) using a 5x widefield (NA 0.16) pre-scan objective and 40x water-immersion objective for the determined organoid area (NA 0.6, working distance 0.62 mm, depth of focus 1,2 µm) using three excitation lasers (405 nm with emission band-pass filter 435/480; ex 488, em 500/550 and ex 561, em 570/630). Nine fields-of-view with 5% overlap were imaged per well using 15 predetermined Z focus planes with laser-based autofocusing. The images were captured with two Andor Zyla sCMOS cameras (16-bit, field of view 650 x 650 μm2, effective xy resolution 0.66 µm). Z-stacks of 15 images were analyzed from a maximum projection using the PerkinElmer Harmony 4.9 software package as described in Supplementary File 1.

### Statistical analysis

Statistical analyses were conducted with R version 4.3.1. The proportional nephrin and ECAD-positive area per total Hoechst-stained area in an organoid were calculated based on the Opera Phenix-acquired images and data files. For the higher throughput approaches 3 and 4, the spheroid organoid diameter was calculated based on the total Hoechst-stained area. Mann–Whitney U test was used for two-group analyses, while the comparison of multiple groups was done using the Kruskal–Wallis test and Spearman’s rank correlation coefficient was used to assess correlation between nephrin and ECAD positive area per organoid. Multiple linear models were fitted using the proportional nephrin- and ECAD-positive area treated as dependent variables and the higher throughput approach (1-4), cell line (HEL24.3 and HEL61.2), Experimental replicates (experiments A and B), and cell number at spheroid initiation (2 000 to 200 000 cells) as explanatory variables. The nephrin and ECAD-proportion values were logit-transformed into a continuous scale using a pseudo-count in order to include zero values in the data. The assumptions of a multiple linear model were assessed with data visualization, which were met although mild skewedness was observed in the Q-Q plot of the residuals. Multiple linear models were fitted and results reported using the logit-transformed data.

## Results

### Generation and characterization of iPSC lines

Differentiation capacity was evaluated in our study utilizing six human iPSC lines, namely HEL11.4, HEL24.3, HEL47.2, HEL61.2, HEL124.1, and HEL124.2. The generation and characterization of HEL61.2 (and its clone HEL61.1), representing a 57-year-old, healthy female donor, and HEL124.1 and HEL124.2, the two clones representing a female newborn GRACILE patient (ZCC11-72), are described in the present study. Other iPSC lines have been published earlier^28–30^.

All here established iPSC clones expressed typical stem cell marker proteins TRA-1-60, OCT3/4, and SSEA3 or SSEA4 (Supplementary Figure S1, data shown for HEL61.2; and Supplementary Figure S2, for HEL124.1 and HEL124.2) as well as endogenous pluripotency-associated genes *NANOG*, *OCT3/4*, *SOX2*, and *TDGF1* (Supplementary Figures S1–2). Absence of the SeV was confirmed by RT-PCR (Supplementary Figures S1–2). All iPSC lines generated in the study were able to differentiate into the three germ layers (Supplementary Figure S1).

The presence of the GRACILE syndrome-causing *BCS1L* mutation, c.A232G (p.S78G)^27^, in HEL124.1 and HEL124.2 was verified by Sanger sequencing (Supplementary Figure S2F). The karyotype of HEL61.2, HEL124.1, and HEL124.2 iPSCs was normal 46 XX (Supplementary Figure S3).

### The GRACILE syndrome molecular phenotype is replicated in patient-derived iPSC-lines differentiated into hepatocyte cells and kidney organoids

In the GRACILE syndrome the incorporation of Rieske iron-sulfur protein (RISP/UQCRFS1) into the mitochondrial respiratory chain complex III is decreased due to the *BCS1L* mutation^36,37^, but the magnitude is highly tissue-specific. Deficiencies in the mitochondrial oxidative phosphorylation pathway tend to manifest poorly in non-differentiated proliferating cells. Loss of RISP and subsequent CIII deficiency has not been detected in cultured fibroblasts from patients with *BCS1L* mutations, whereas it is detectable in liver samples^36,38^. Therefore, the molecular phenotype of the novel GRACILE iPSC lines, HEL124.1 and HEL124.2, was assessed with Western blot analysis of mitochondrial respiratory chain proteins in iPSC-derived definitive endodermal cells differentiating towards hepatocyte lineage cells^34^ (Supplementary Figure S4A–B) and kidney organoids^4^ (Supplementary Figure S4AC–D). Original uncropped, unadjusted Western blot images are shown in Supplementary File 2.

The analysis of the definitive endodermal cells with two healthy donor (HEL24.3 and HEL61.2) and both GRACILE iPSC lines (HEL124.1 and HEL124.2) showed adequate endodermal differentiation capacity and confirmed that the expression of RISP and BCS1L were decreased in the GRACILE iPSC-derived hepatocyte lineage cells (Supplementary Figure S4B). Moreover, kidney organoids were generated with the previously published APEL/air-medium interface protocol^4^ (Figure 1A) with the two healthy donor iPSC lines and two batches of GRACILE iPSC line HEL124.2. The expression of both BCS1L and RISP were also low in the GRACILE kidney organoids as compared to that in the healthy donor organoids (Supplementary Figure S4D). All other analyzed mitochondrial proteins (NDUFA9 for complex I, SDHA for complex II, Core1 for complex III, and COX1 for complex IV) were unaffected (Supplementary Figure S4A-D), indicating that the loss of RISP is replicated in the iPSC-derived hepatocyte and kidney tissue models of the GRACILE syndrome.

### The APEL/air-medium interface protocol shows variability in kidney organoid structure development across cell lines

Success rates and replicability of the original APEL/air-medium interface kidney organoid protocol^4^ (Figure 1A) were further evaluated across seven stem cell lines, which were not preselected based on their capacity to differentiate towards kidney cells. The success rate was determined based on the appearance of intrinsic structures seen under the light microscope and subsequently confirmed by a fluorescence microscopy analysis of markers for maturing kidney cells.

The overall success rate by day 20 was encouraging when tested with six iPSC lines and one hESC line (Figure 2, Table 1). However, the success rates varied between the cell lines with some producing organoids from experiment to experiment, while others showed more variable outcomes, and the healthy-donor line HEL47.2 and GRACILE line HEL124.1 failed to produce kidney organoids (Figure 2, Table 1). Interestingly, while GRACILE cell line clone HEL124.2 showed good success rates overall in our study, in four initial experiments the organoids showed tubular degradation before reaching day 20 (Supplementary Figure S5).

**Figure 2.**
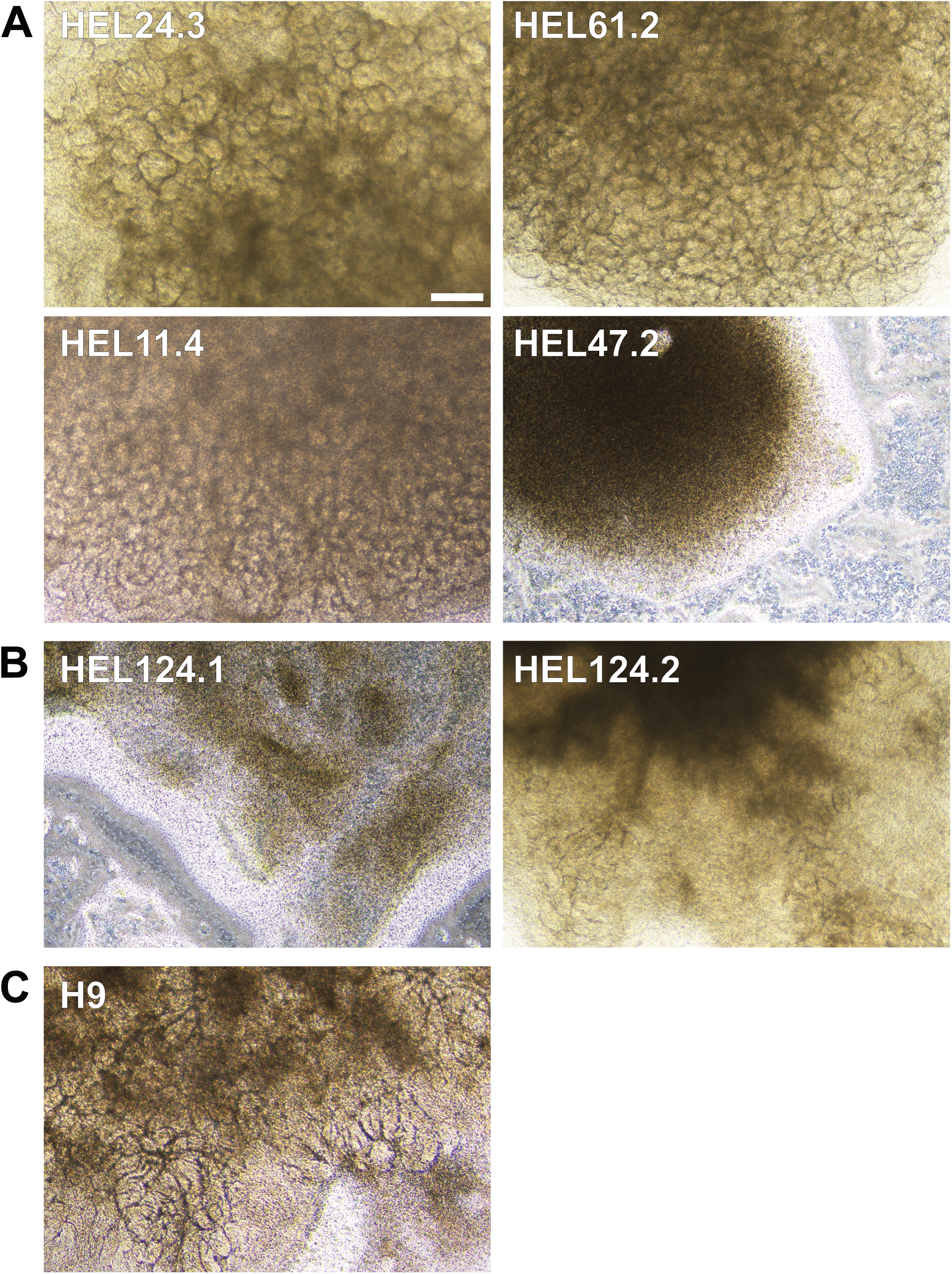
Representative light microscopy images of kidney organoids generated with seven stem cell lines to test success rates and reproducibility of the APEL/air-medium interface protocol. Six hiPSC lines, complemented by one hESC line H9, were tested with the original APEL/air-medium interface protocol for kidney organoid differentiation for 20 days. Representative light microscopy images of kidney organoids generated with (A) healthy-donor iPSC lines HEL24.3, HEL61.2, HEL11.4, and HEL47.2 (B) two GRACILE-syndrome patient-derived clonal iPSC lines HEL124.1 and HEL124.2 (C) a hESC line H9. (A-C) HEL47.2 and HEL124.1 did not produce kidney organoids while other tested cell lines generated organoids with intrinsic structures. Scale bar 200 µm.

**Table 1.** Success rate of kidney organoid differentiation utilizing the original APEL/air-medium interface protocol.

| Cell lines | All |  | <50% cell plating density at D0 |  |  | > 50% cell plating density at D0 |  |  |
| --- | --- | --- | --- | --- | --- | --- | --- | --- |
|  | Success rate (%)<sup>1</sup> | Number of samples<sup>2</sup> | Success rate (%)<sup>1</sup> | Number of samples<sup>2</sup> | Number of experiments | Success rate (%)<sup>1</sup> | Number of samples<sup>2</sup> | Number of experiments |
| **hiPSC** |  |  |  |  |  |  |  |  |
| HEL24.3 | **69** | 35 | 95 | 21 | 6 | 29 | 14 | 4 |
| HEL47.2 | **0** | 3 | ND | ND | NA | 0 | 3 | 2 |
| HEL61.2 | **100** | 47 | 100 | 29 | 7 | 100 | 18 | 5 |
| HEL124.1 | **0** | 5 | 0 | 1 | 1 | 0 | 4 | 2 |
| HEL124.2 | **96** | 45 | 97 | 30 | 9 | 93 | 15 | 5 |
| HEL11.4 | **100** | 3 | 100 | 3 | 1 | ND | ND | NA |
| **hESC** |  |  |  |  |  |  |  |  |
| H9 | **92** | 12 | 100 | 6 | 2 | 83 | 6 | 2 |

Note:<sup>1</sup>Evaluated based on the appearance of intrinsic structures in light and immunofluorescence microscope. <sup>2</sup>Total number of samples, including different experiments. The same or subsequent batches of each iPSC line were utilized. For HEL61.2, HEL124.1, and HEL124.2, both low and high passage iPSC batches were additionally tested. Abbreviations: D0, day 0 of the differentiation; hiPSC, human induced pluripotent stem cell; hESC, human embryonic stem cell; NA, not applicable; ND, not determined

Immunostaining with kidney cell markers demonstrated structures of nephrin-positive podocyte cells, ECAD-positive maturing tubular epithelial cells, and Lotus tetragonolobus lectin (LTL)-positive proximal tubule cells (representative images shown for healthy-donor cell lines HEL24.3, HEL61.2, and GRACILE cell line HEL124.2 in Figure 3 and Supplementary Figure S6). Experiment-to-experiment variation was observed, in the number of nephrin-positive glomerular structures in particular, yet the overall appearance of tubular structures at the end of the differentiation was primarily cell line-dependent. Compared with the other tested cell lines, the healthy donor iPSC line HEL61.2 produced organoids with seemingly more complex tubular structures (Figure 3, Supplementary Figure S6).

**Figure 3.**
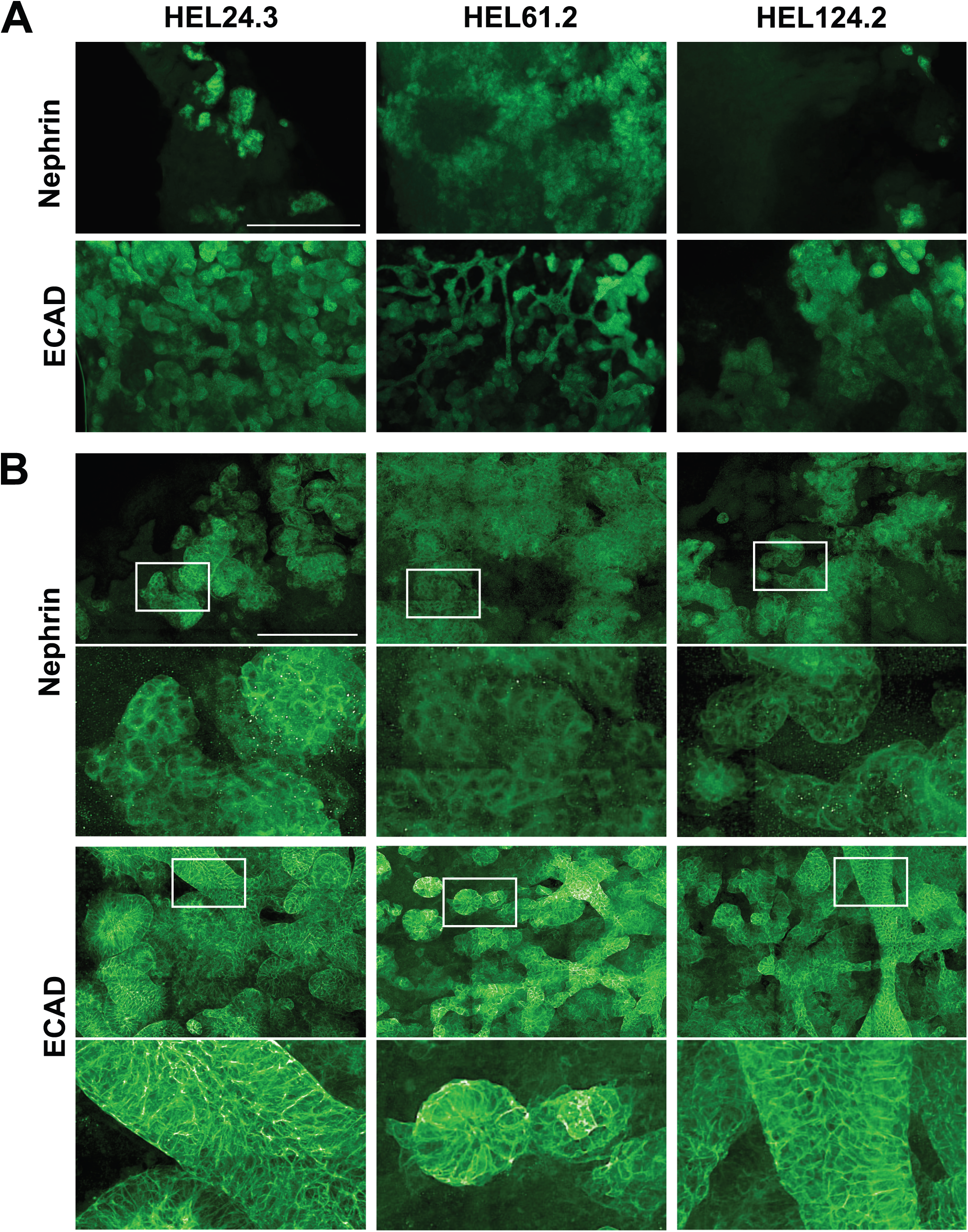
Morphology of glomerular and tubular structures in iPSC-derived kidney organoids generated with the original APEL/air-medium interface protocol. Representative immunofluorescence images of the maturing glomerular structures, stained for nephrin, and maturing tubular structures, stained for epithelial cadherin (ECAD) in healthy-donor iPSC lines HEL24.3 and HEL61.2, and the GRACILE iPSC line HEL124.2, images acquired with (A) widefield fluorescence microscopy, scale bar 500µm, and (B) the Opera Phenix spinning disk confocal microscope, scale bar 200µm.

### Optimization of the APEL/air-medium interface protocol towards higher throughput shows association between culture conditions and kidney organoid structure development

Since the original APEL/air-medium interface protocol is laborious and the organoids need to be manually processed one by one in relevant steps, the protocol was modified towards higher throughput. Altogether four modified approaches with different culture systems and amounts of cells were tested for their ability to support the differentiation while enabling simultaneous processing of a plateful of samples (24–96 wells) (Figure 1B). Comparable to the original protocol (Figure 1A), two of the approaches were started with monolayer cultures (Figure 1B, approach 2 and 3). However, all approaches utilized 96-well-plate spheroid suspension cultures (one spheroid per well) for the generation of small, 24- and 96-well-compatible spheroids (Figure 1B, approaches 1–4). The spheroids were either placed on membranes at the air-medium interface for final differentiation (24-well utilized in the optimization stage; Figure 1B, approaches 1 and 2) or were differentiated further as spheroids (Figure 1B, approaches 3 and 4).

The approaches were simultaneously tested with two healthy-donor iPSC lines, HEL24.3 and HEL61.2, and in two separate experiments (experiment A and B). Since spheroid size may affect the number of differentiated structures^14^, cultures were started with a variable number of cells, with approaches 1 and 4 spheroids formed using 2 000 to 50 000 cells at day 0 and approaches 2 and 3 spheroids formed using 100 000 or 200 000 cells at day 7. To image and quantify the proportional amount of immunostained kidney structures, a pipeline for Opera Phenix high-content screening analysis of the organoids was developed (Supplementary File 1). For the screening we selected podocyte-specific nephrin as a marker for glomerular structures, and ECAD as a marker for tubular structures, as it should recognize adequate number or maturing proximal/distal tubule epithelial cells, considering that the kidney organoids generated here have not reached full maturity^4,35^.

Opera Phenix analysis demonstrated that all tested higher throughput approaches allowed the formation of structures positive for the podocyte and tubular markers (Figure 4 and Figure 5). Experiment-to-experiment variation in structure development was observed, with the approaches utilizing spheroid cultures in particular (Figure 4, compare the approaches 3 and 4 between A and B). Moreover, approaches 3 and 4 produced spheroid organoids of uniform size by day 20 of culture with good replicability across cell lines, however more experiment-to-experiment variation was seen for approach 3 (Supplementary Figure S7A). With structure development there were challenges in replicability for all included samples (Figure 4, compare A and B for both nephrin and ECAD in HEL61.2 with approach 1 and 10 000 cells). Nevertheless, there was a trend indicating that the utilization of air-medium interface at the end of the differentiation had overall more potential in producing kidney structures compared with the approaches continuing with spheroid cultures to the end of the differentiation (Figure 4, showing results of two independent experiments, Supplementary Figure S7B showing pooled data from Figure 4; and Figure 5 showing representative Opera Phenix immunofluorescence images for the comparison between the approach 4 and 1). On the other hand, the spheroid approaches 3 and 4 allowed for the generation of a larger number of organoids providing more statistical power (Supplementary Table S3). Accordingly, the most statistically significant correlation between nephrin and ECAD-positive structure development per organoid was seen for the spheroid approach 4, although the other approaches supported the overall structure development better (Supplementary Figure S7C).

**Figure 4.**
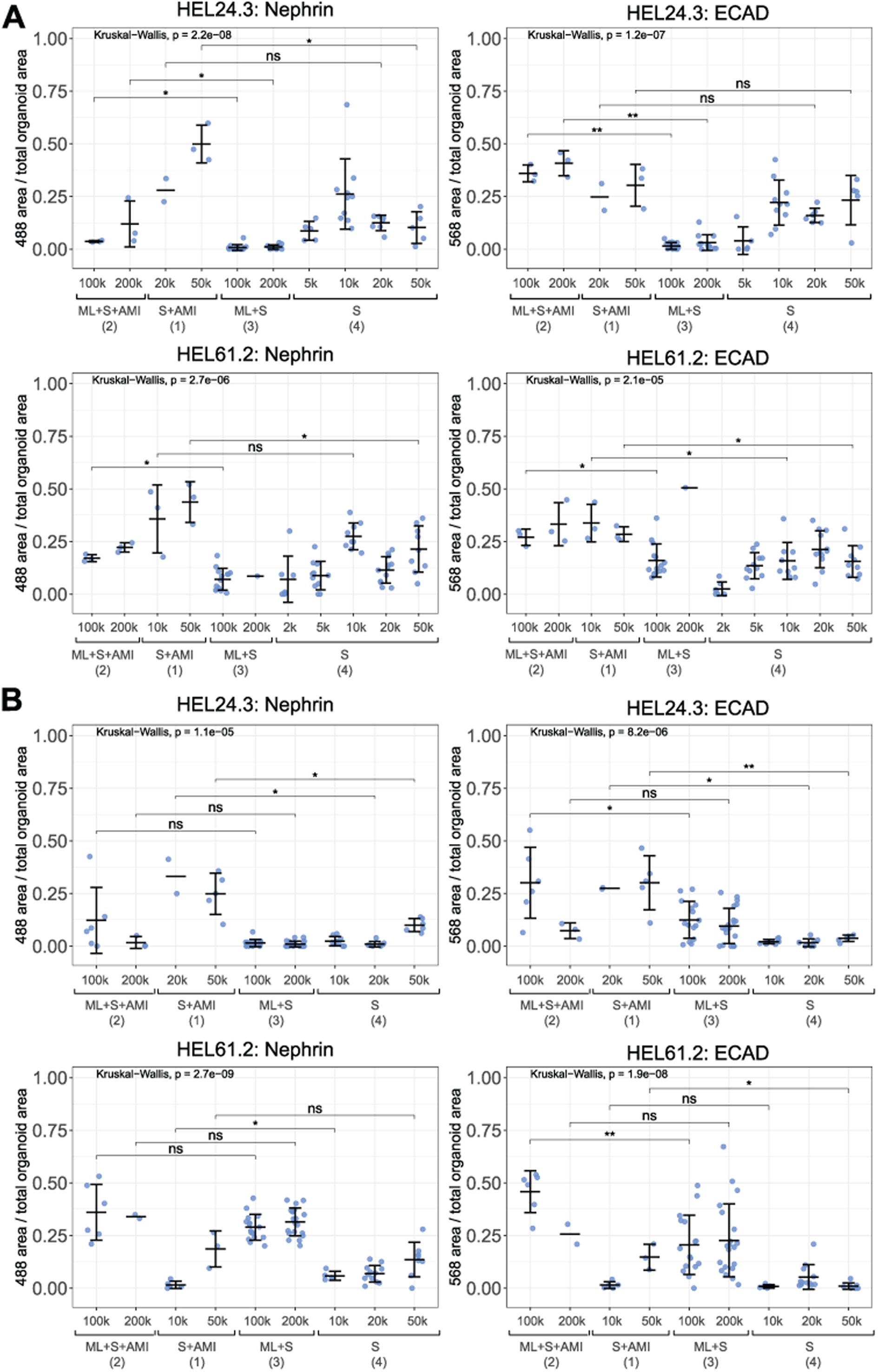
Opera Phenix high-content screening and quantification of nephrin and ECAD-positive maturing structures in kidney organoids generated with the higher throughput approaches. Four modified high-throughput approaches consisting of different culture systems (monolayer culture, ML, spheroid suspension culture, S, and/or air-medium interface, AMI) and variable amounts of cells at the initiation of spheroids, from 2 000 (2k) to 200 000 (200k) cells, were tested. Two iPSC lines were utilized (HEL24.3 and HEL61.2) in two different experiments (A and B). Proportion of nephrin and ECAD-positive area per organoid (imaged with Opera Phenix) is shown with a dot (mean ± s.d. for each condition indicated with black crossbars). Unadjusted significance levels are shown (Mann-Whitney U test *: p < 0.05, **: p < 0.01, ns: non-significant). Number of replicate samples per condition provided in Supplementary Table S3.

**Figure 5.**
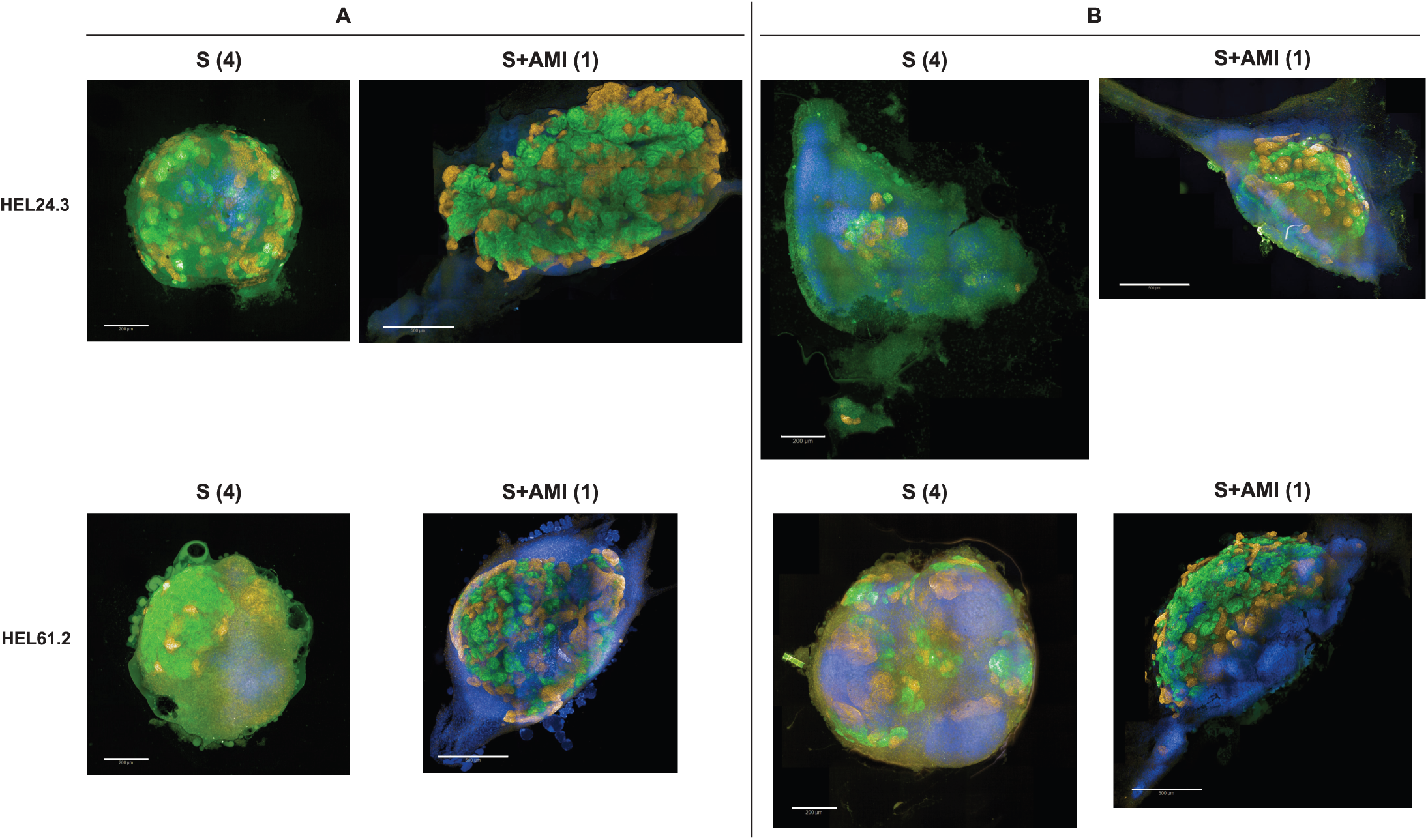
Representative Opera Phenix immunofluorescence images of nephrin and ECAD-positive maturing structures in kidney organoids generated with the higher throughput approaches. Kidney organoids generated with the high-throughput approaches 1 (S+AMI, a spheroid suspension culture followed by air-medium interface) and 4 (S, a spheroid suspension culture). Images represent two iPSC lines HEL24.3 and HEL61.2 and two separate experiments (A and B) with 50 000 cells utilized for the initiation of spheroids. Organoids were stained for nephrin (green) and ECAD (red) for Opera Phenix high-content screening. Scale bar 200 µm. (S) and 500 µm (S+AMI).

The effects of the high-throughput approaches and culture conditions on the observed structure variability were further assessed with multiple linear models, using the high-throughput approach, cell line, experimental replication and cell number at spheroid initiation as explanatory variables (Tables 2-3). Since approaches S+AMI (1) and S (4) used initial cell number ranging from 2 000 to 50 000 for spheroid formation at day 0, they were analyzed as a separate subgroup from approaches ML+S+AMI (2) and ML+S (3), which used initial cell number ranging from 100 000 to 200 000 for spheroid formation at day 7. For subgrouped approaches 1 and 4, cell line showed no significant effect on structure formation, while cell number at spheroid initiation and the experiment played a significant role. The model for approaches 1 and 4 explained 35% of the variability in the logit-transformed proportion of nephrin-positive area and 48% of the variability in the logit-transformed proportion of ECAD-positive area. On the contrary, for subgroup of approaches 2 and 3, cell line, and experiment had a significant impact, while cell number at spheroid initiation had no significant impact on structure formation. The model of subgrouped approaches 2 and 3 explained 76% of the variability in the logit-transformed proportion of nephrin-positive area and 44% of the variability in the logit-transformed proportion of ECAD-positive area. Both subgroup analyses indicated that the approaches utilizing air-medium interface at the end of the protocol, S+AMI (1) and ML+S+AMI (2), increased structure formation compared to the approaches, ML+S (3) and S (4), utilizing spheroid culture to the end. The approach 1 (spheroid suspension culture followed by AMI), the most potential approach in terms of simplicity and ability to support the differentiation, was also successfully adapted to 96-well level with robotics-friendly array of 96-well membrane inserts (Supplementary Figure S8).

**Table 2.**
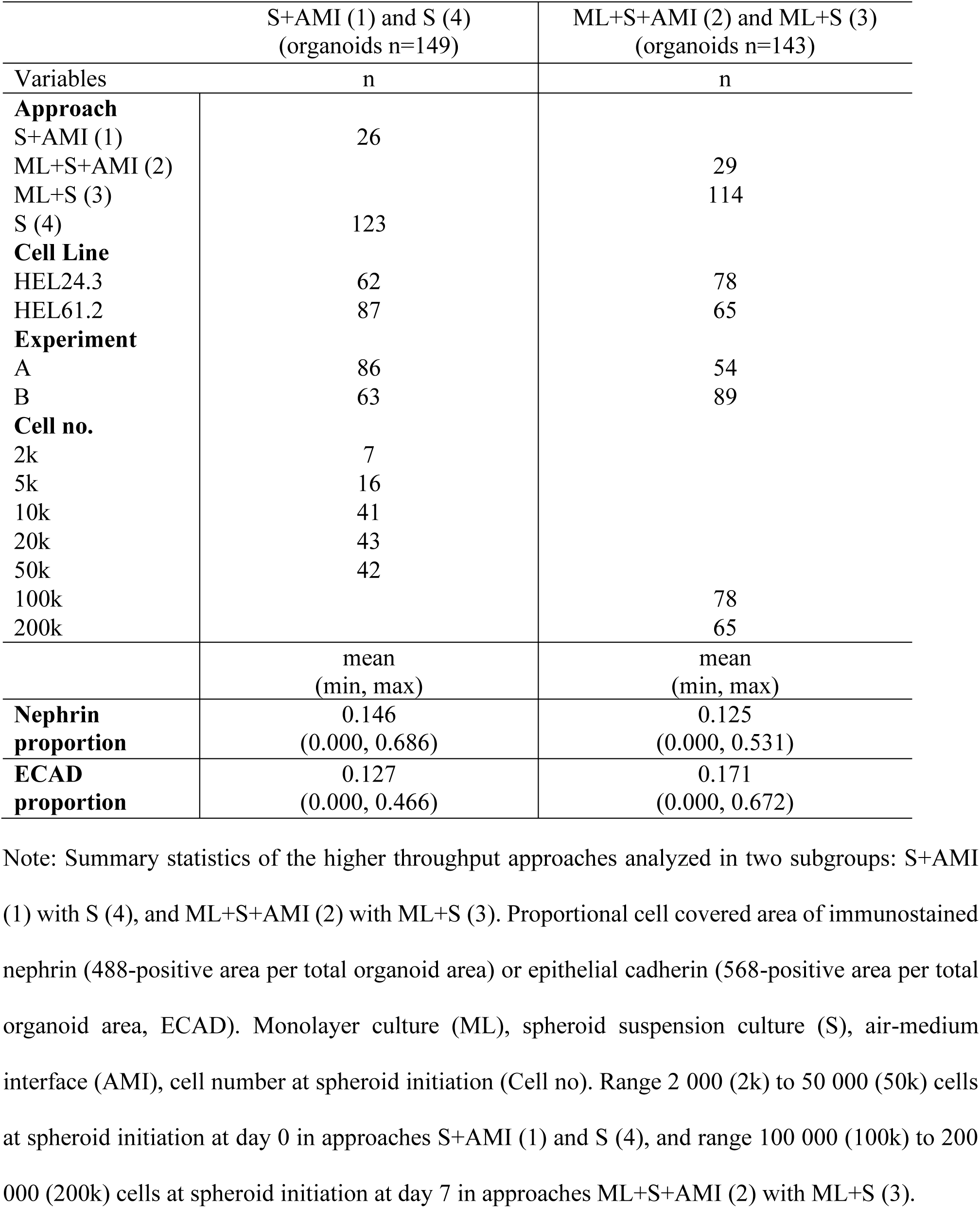
Summary statistics of kidney organoids differentiated with the higher throughput approaches and imaged with Opera Phenix.

**Table 3.** Association between higher throughput culture conditions and kidney organoid glomerular (nephrin) and tubular (ECAD) structures assessed with multiple linear models.

|  | S+AMI (1) and S (4)<br>(organoids n=149) |  | ML+S+AMI (2) and ML+S (3)<br>(organoids n=143) |  |
| --- | --- | --- | --- | --- |
|  | Nephrin<br>488 | ECAD<br>568 | Nephrin<br>488 | ECAD<br>568 |
| Approach:<br>S+AMI (1) | 0.921***<br>(0.223) | 1.120***<br>(0.209) |  |  |
| Approach:<br>ML+S+AMI (2) |  |  | 0.893***<br>(0.158) | 1.481***<br>(0.215) |
| Cell Line:<br>HEL61.2 | 0.014<br>(0.167) | -0.265<br>(0.156) | 2.486***<br>(0.129) | 1.206***<br>(0.176) |
| Experiment: B | -1.223***<br>(0.170) | -1.677***<br>(0.160) | 0.691***<br>(0.134) | 0.577**<br>(0.182) |
| Cell no. | 0.023***<br>(0.005) | 0.015**<br>(0.005) | 0.001<br>(0.001) | -0.001<br>(0.002) |
| $r^2$ | 0.371 | 0.489 | 0.771 | 0.451 |
| Adjusted $r^2$ | 0.353 | 0.475 | 0.764 | 0.435 |

Note: Higher throughput approaches analyzed in two subgroups: S+AMI (1) with S (4), and ML+S+AMI (2) with ML+S (3). Multiple linear models fitted with logit-transformed proportions of immunostained nephrin (488-positive area per total organoid area) or epithelial cadherin (568-positive area per total organoid area, ECAD). Coefficients of the logit-transformed proportions are shown with standard error in brackets for the variables. Significance levels \*\*: $p < 0.01$ , \*\*\*: $p < 0.001$ . Monolayer culture (ML), spheroid suspension culture (S), air-medium interface (AMI), cell number at spheroid initiation (Cell no.)

## Discussion

Human iPSC-derived kidney organoids represent an alternative strategy to generate tissue models for the studies of genetic diseases, disease mechanisms, and drug screening^5^. Along with the efforts to enhance the maturation, glomerular vascularization, and the functionality of the kidney cells^39–41^, it is also crucial to report on line-to-line and experiment-to-experiment variations that may play a significant role, especially in long multistep differentiations. These variations are a known challenge in the iPSC field, but remain scarce in the kidney organoid literature, for negative results in particular^20,42^. Previous studies have utilized transcriptomics for the testing of a specific protocol (the protocol of Takasato et al.) and have shown variation in the maturation and abundance of kidney and non-kidney cells of the organoids produced from the same clone but at a different time^43^.

In addition to the experimental variation, the protocol and cell line of choice may play a significant role in the transferability of the methods across laboratories. Available kidney organoid protocols differ in terms of basal medium, targeted signaling pathways, the concentrations and timings of each chemical induction, culture systems and vessel format, and cell lines used in the establishment of the protocol. Moreover, each patient-derived iPSC line and its subclones need to be extensively tested to select optimal conditions for e.g., functional analyses, which further increases the methodological complexity in this field. The APEL/air-medium interface protocol^4^ by Takasato et al. showed variable but generally positive success rates in our study. Notably, by reporting positive and negative rates of success with cell lines not preselected based on their differentiation capacity, we demonstrate that iPSC-derived kidney organoids can be generated with rather high success rates, but the overall success and structure development can be highly cell line dependent. The negative success rates observed in the present study, however, do not exclude the possibility that the tested cell lines may show different outcomes with protocols and induction methods not tested in the study. Previous studies have compared kidney organoid protocol outcomes, e.g. by reporting transcriptional differences in cell type ratios and maturation state between the organoids derived using the protocols of Takasato et al. and Morizane and Bonventre^21^. Moreover, a modified Takasato protocol has been utilized in the validation of the development of nephron-like structures generated in the established adherent high-throughput differentiation^11^. The present study highlights the importance of high-throughput applications of morphological analyses, along with the frequently applied omics approaches, as immunostaining-based analyses remain indispensable in mapping the structural orchestration of organoids.

Although different iPSC lines may have variable outcomes in kidney organoid differentiation, air-medium interface was here found to have in general a positive impact on the production of glomerular and tubular structures, and based on the present study and the data of others, it can be adapted to higher throughput with the possibility to control the size of an organoid^18,19,44^. High-throughput applications, e.g., functional analyses and drug-based screenings, will need a large number of uniform-sized organoids. These requirements may be difficult to obtain with co-cultures of more than one organoid. Published high-throughput kidney organoid protocols have utilized co-cultures or suspension cultures of multiple organoids ^11,14^, which may not be optimal for reliable screening and statistical analyses, as technical variation can significantly affect organoid size and maturation^45^. Indeed, bioprinting has been successfully utilized in the placement of the cells at air-medium interface to increase throughput of the APEL/air-medium interface protocol and to control the size of the organoids^44^. In the present study, a spheroid suspension culture step was incorporated in the protocol to reduce and control the size of the organoids, which were cultured in individual wells up to 96-well level at air-medium interface. Thereby our method can enable better statistical management of technical and biological sources of variation, such as the effects of cell lines, initial cell populations, or experimental replication. Although limited by efficiency when compared with bioprinting, our microplate-based approach can serve as an accessible option for kidney organoid screening studies in laboratories where cell printing technology is not available.

Our results also showed that kidney organoids represent a promising tissue model for the study of the GRCILE syndrome, as the molecular Rieske iron-sulfur protein (RISP/UQCRFS1) phenotype of the disease was replicated in iPSC-derived kidney cells, in a cell-autonomous manner. The GRACILE syndrome is caused by a homozygous missense mutation in *BCS1L* (c.A232G, p.S78G), which encodes a translocase required for the incorporation of RISP into mitochondrial Complex III^27^. Proximal tubulopathy is an early postnatal feature of the disease, causing generalized Fanconi type aminoaciduria with increased excretion of lactate, glucose, pyruvate, and phosphate^26^. The syndrome has previously been studied in human tissue samples and in a knock-in mouse model with the human mutation^36–38,46,47^. The present study is the first where GRACILE patient-derived iPSCs and respective tissue models have been generated, but full functional characterization was not within the scope of this study. Of note, only one out of the two patient-derived clonal GRACILE iPSC lines produced kidney organoids, with some experiment-to-experiment variation in viability. Future studies with additional patient cell lines and isogenic controls are needed as our preliminary attempts to utilize the CRISPR/Cas9 approach in the generation of genome-edited, isogenic control cell line for GRACILE HEL124.2 were not successful.

Our study highlights the potential of iPSC-derived tissue organoids for the modelling of genetic and mitochondrial diseases but also underline the importance of extensive testing of available iPSC lines and methodology before proceeding to large-scale functional analyses and comparison of data obtained by different laboratories. Our high-throughput approach for the production and morphological analysis of kidney organoids may enable more robust distinction between technical and biological sources of cell type variability. Moreover, the method can help combat the issue of size variability and is likely easily adaptable by other laboratories. Overall, the present study promotes the validation, transparency, and transferability of kidney organoid technologies across laboratories, hopefully serving as a reference for similar studies in the future.

## Acknowledgments

The authors thank BSCC supported by HiLIFE and Biocenter Finland for iPSC services and support on hepatocyte differentiation. The authors acknowledge the FIMM High Content Imaging and Analysis Unit; FIMM-HCA (FIMM, HiLIFE, UH; Biocenter Finland & Euro-Biomaging) for provided imaging services.

## Authorship contribution statement

Conceptualization: KUR, RT, JK, ML, VF, PHG. Data Curation: KUR, AP, RT. Formal Analysis: KUR, AP, RT. Funding Acquisition VF, PHG, ML, KUR, AP. Investigation: KUR, AP, RT, AH. Methodology: KUR, RT, AH. Project Administration: KUR, ML. Resources: SL, ML, VF, PHG. Software: Not applicable. Supervision: ML, VF, JK, PHG. Validation: KUR, AP, AH, RT, SL. Visualization: KUR, AP, AH, RT. Writing – Original Draft Preparation: KUR, AP. Writing – Review & Editing: All authors.

## Data availability statement

The informed consent signed by the donors/guardians does not cover publication of original patient DNA sequencing data, all other original data will be made available upon reasonable request from the FinnDiane Study Group/the corresponding author for non-commercial research purposes.

