## Supplemental File S2 for "Parsing glomerular and tubular structure variability in high-throughput kidney organoid culture"

**Supplementary File 2.**
Original uncropped and unadjusted blot images underlying all Western blot results reported in the present study. The original blots were cut horizontally for simultaneous probing of different proteins per blot (>50 kDa, 37 – 50 kDa, and < 37 kDa) with well-established antibodies.

<u>The sample order from left to right in Supplementary Figure S4, panels A and B (definitive endoderm samples):</u>

1. HEL24.3 definitive endoderm (DE)
2. HEL124.1
3. HEL124.2
4. HEL24.3
5. HEL124.1
6. HEL124.2
7. HEL24.3
8. HEL61.2*
9. HEL124.1
10. HEL124.2

Supplementary Figure S4, A, SDHA [the upper blot of the three blots in right, which were scanned together (one blot was negative)]

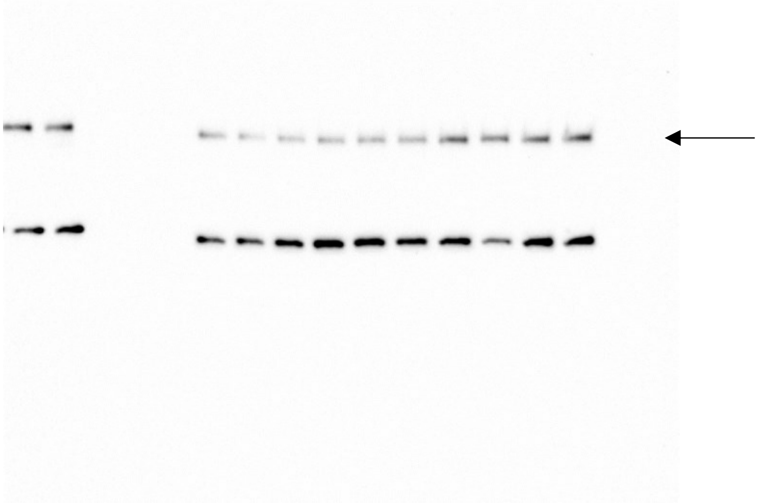

Supplementary Figure S4, A, b-actin [the middle blot of the three blots in right, which were scanned together (one blot was negative)]

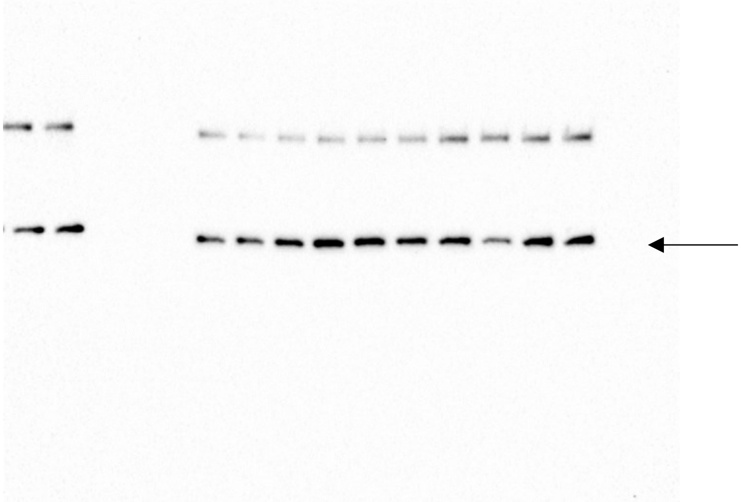

Supplementary Figure S4, A, porin (the lower blot of the two blots which were scanned together)

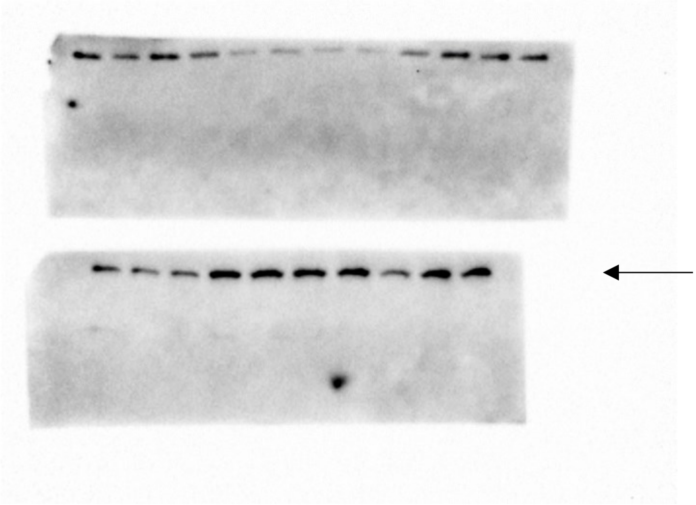

Supplementary Figure S4, B, Core 1 (the lower blot of the two blots which were scanned together)

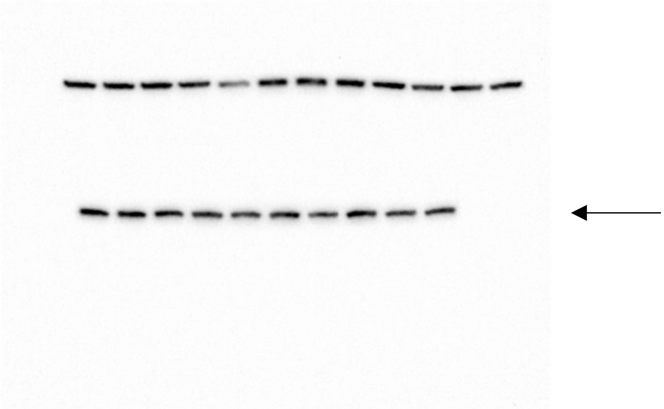

Supplementary Figure S4, B, BCS1L (the lower blot, with a standard lane, of the two blots which were scanned together)

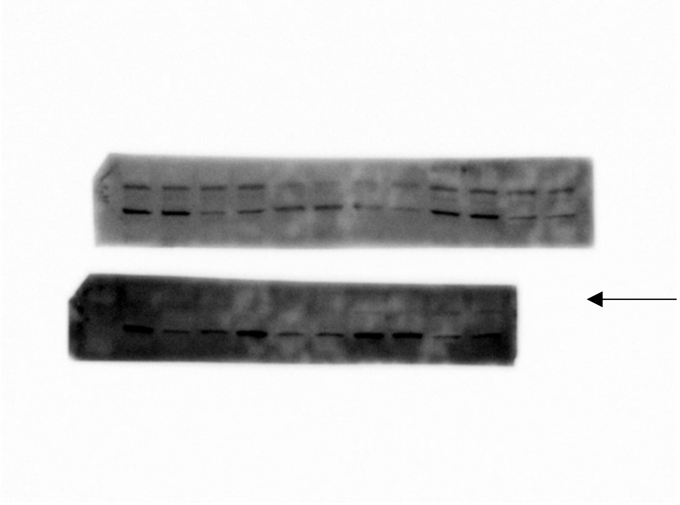

Supplementary Figure S4, B, porin (the lower blot of the two blots which were scanned together)

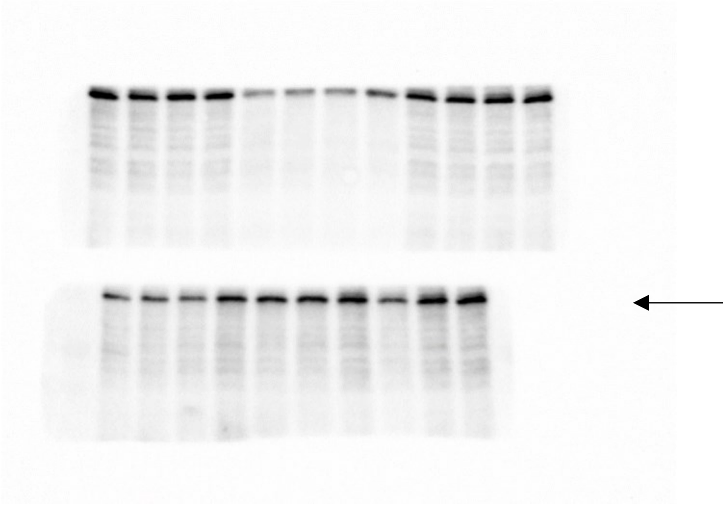

Supplementary Figure S4, B, RISP (the lowest blot of the three blots in left, which were scanned together)

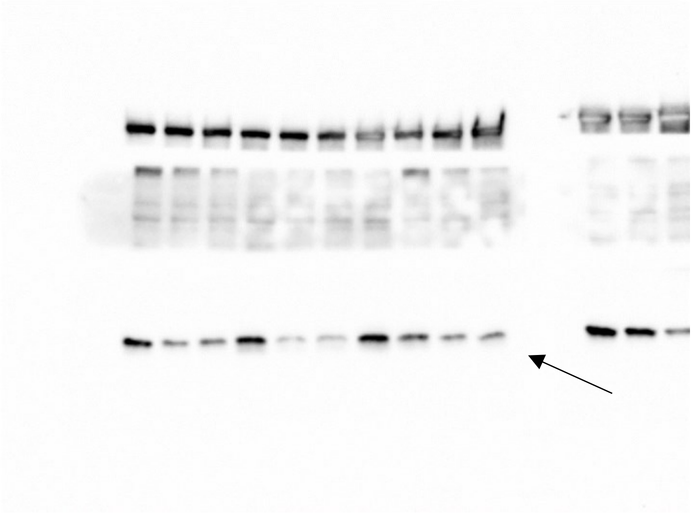

<u>The sample order from left to right in Supplementary Figure S4, panels C and D (kidney organoid samples)</u> is comparable to the order in Fig.6, except that in the original uncropper blots, there were two additional staining controls at left, which were not included in the analysis. The thick arrow indicates the first sample included in the analysis (i.e., 1. HEL24.3).

1. HEL24.3
2. HEL61.2
3. HEL124.2
4. HEL124.2
5. HEL24.3
6. HEL24.3
7. HEL61.2
8. HEL61.2
9. HEL124.2
10. HEL124.2
11. HEL124.2

Supplementary Figure S4, C, SDHA and Supplementary Figure S4, C, b-actin (the two stainings scanned at the same time)

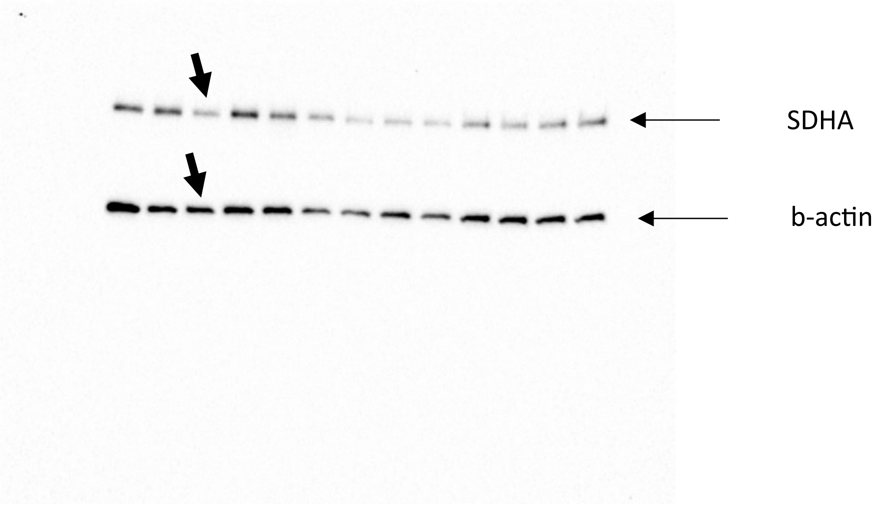


Supplementary Figure S4, C, porin

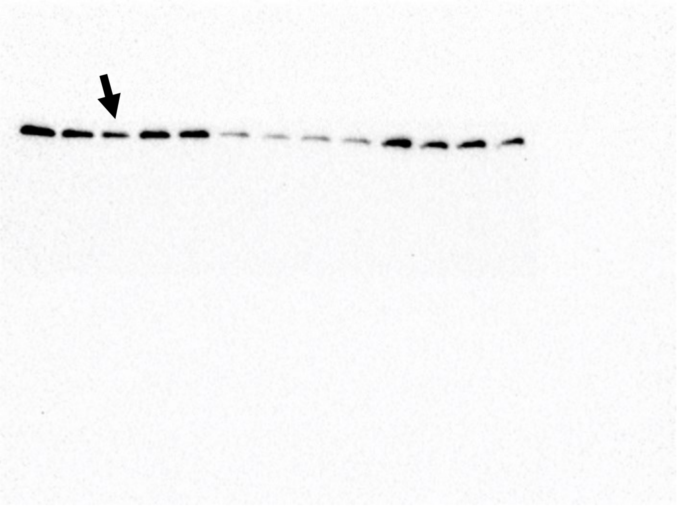

Supplementary Figure S4, D, Core 1 (the middle blot of the three blots which were scanned together)

- This particular blot was first stained for BCS1L (indicated by a star) and then for Core 1 (stripped and restained)

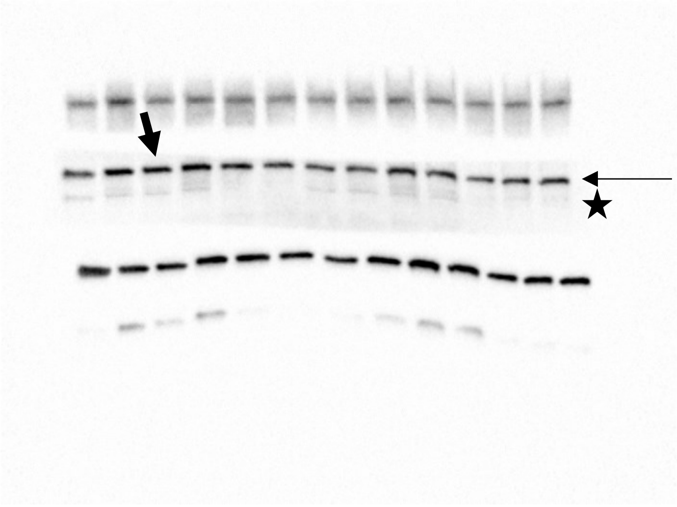

Supplementary Figure S4, D, BCS1L

- This blot was first stained for Core 1 (indicated by a star) and then for BCS1L without stripping between the two stainings to avoid the loss of BCS1L with a faint signal

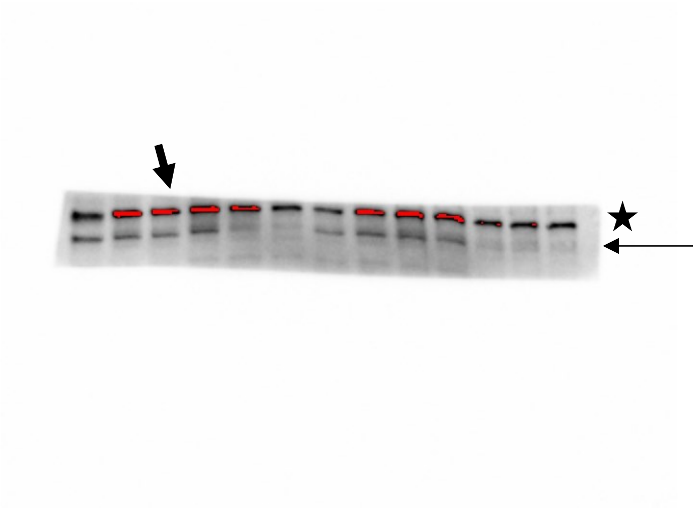

Supplementary Figure S4, D, porin (the lowest blot of the three blots which were scanned together)

- This blot was first stained for RISP (indicated by a star) and then for porin (stripped and restained)

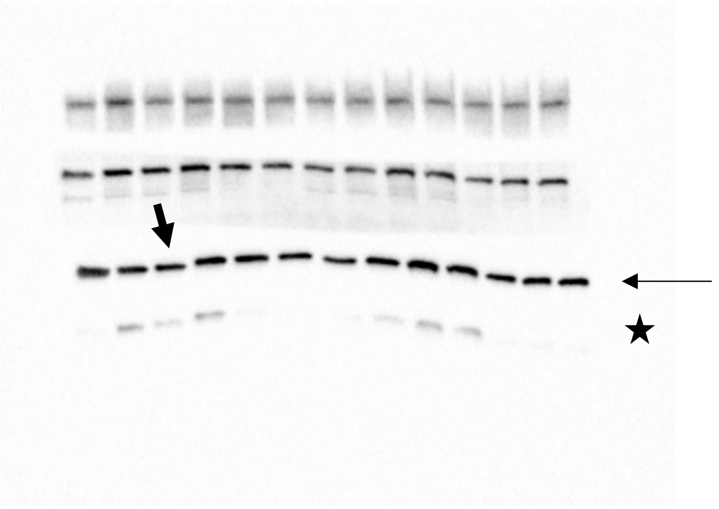

Supplementary Figure S4, D, RISP (the lower blot of the two blots which were scanned together)

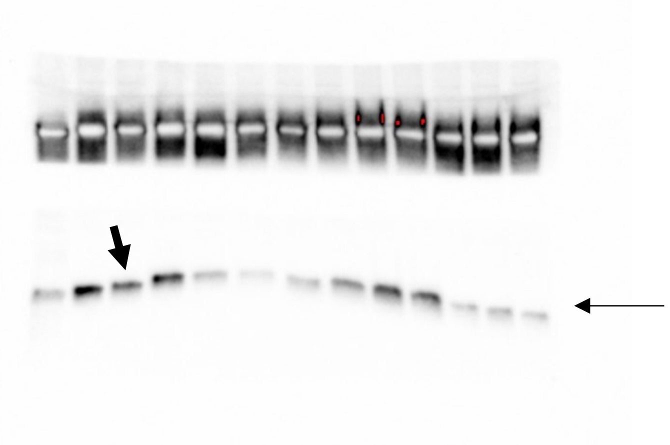

Supplementary Figure S4, D, SDHA (the lower blot of the two blots which were scanned together)

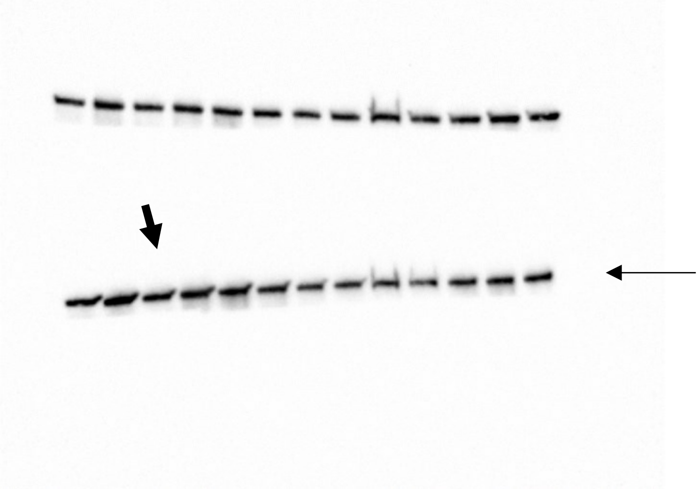

Supplementary Figure S4, D, NDUFA9 (the lower blot of the two blots which were scanned together)

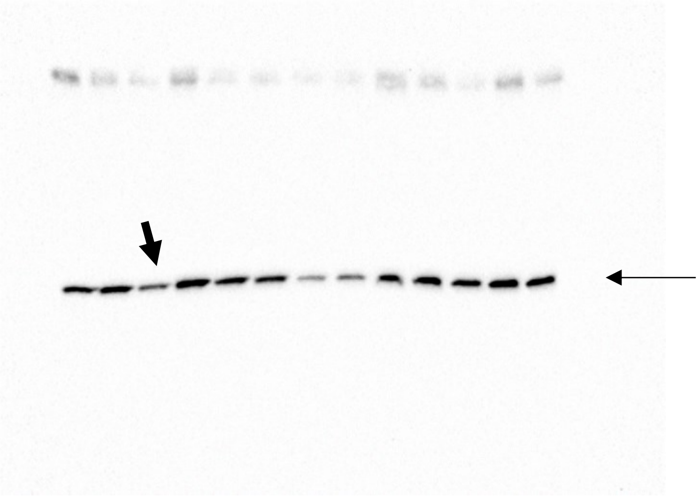

Supplementary Figure S4, D, SDHA (the upper blot of the two blots which were scanned together)

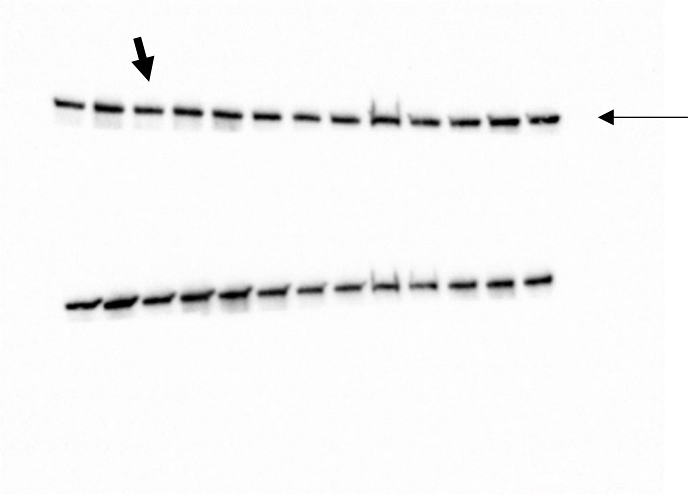

Supplementary Figure S4, D, Cox-1

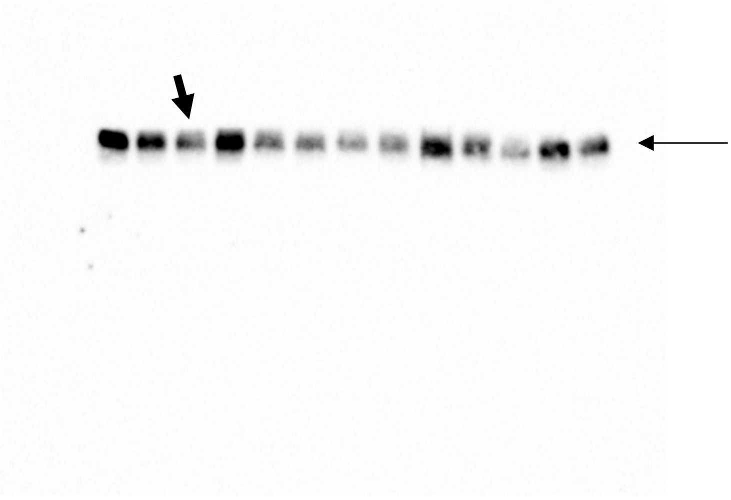
