## Supplementary material for "Parsing glomerular and tubular structure variability in high-throughput kidney organoid culture"

### Supplementary Figures and Tables

#### Supplementary Figures

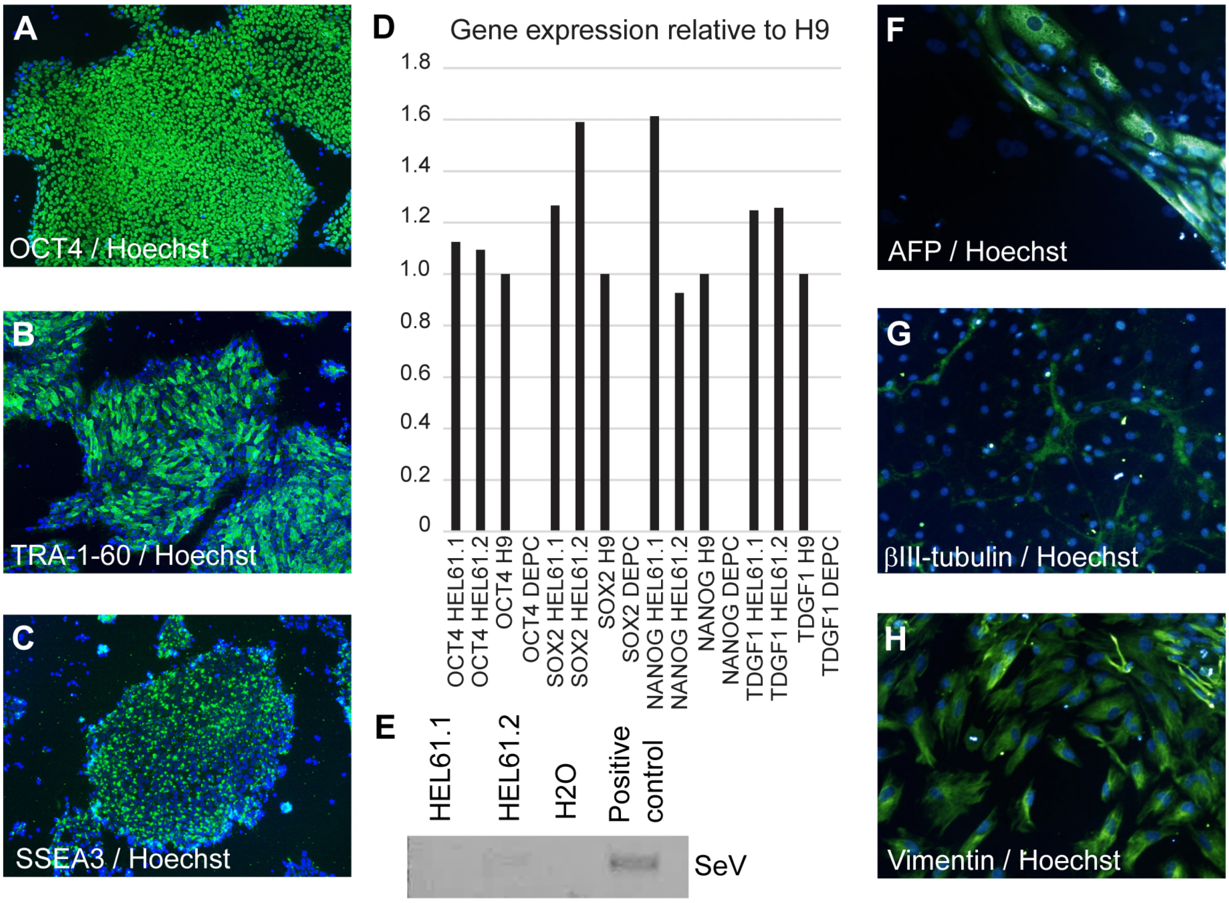


**Supplementary Figure S1. Characterisation of iPSC clones HEL61.2 and HEL61.1 from a 57-year-old female healthy donor.** (A–C) HEL61.2 and HEL61.1 had typical colony morphology and were positive for the nuclear/plasma membrane stem cell markers OCT4, TRA-1-60, and SSEA3 (representative immunofluorescence microscopy images of HEL61.2 with the nuclear Hoechst staining in blue). (D) HEL61.2 and HEL61.1 expressed endogenous stem cell marker genes, *OCT3/4*, *SOX2*, *NANOG*, and *TDGF1* verified by RT-qPCR (hES cell line H9 used as a positive control). (E) Viral expression monitored at passage eight by RT-PCR using primers specific for the Sendai virus (SeV). Full clearance was observed from passage 13 onwards. (F–H) Pluripotency was verified by staining embryoid body-derived cultures for the markers of the three germ layers, α-fetoprotein (AFP, endoderm), β-III tubulin (ectoderm), and vimentin (mesoderm; representative immunofluorescence microscopy images of HEL61.2).

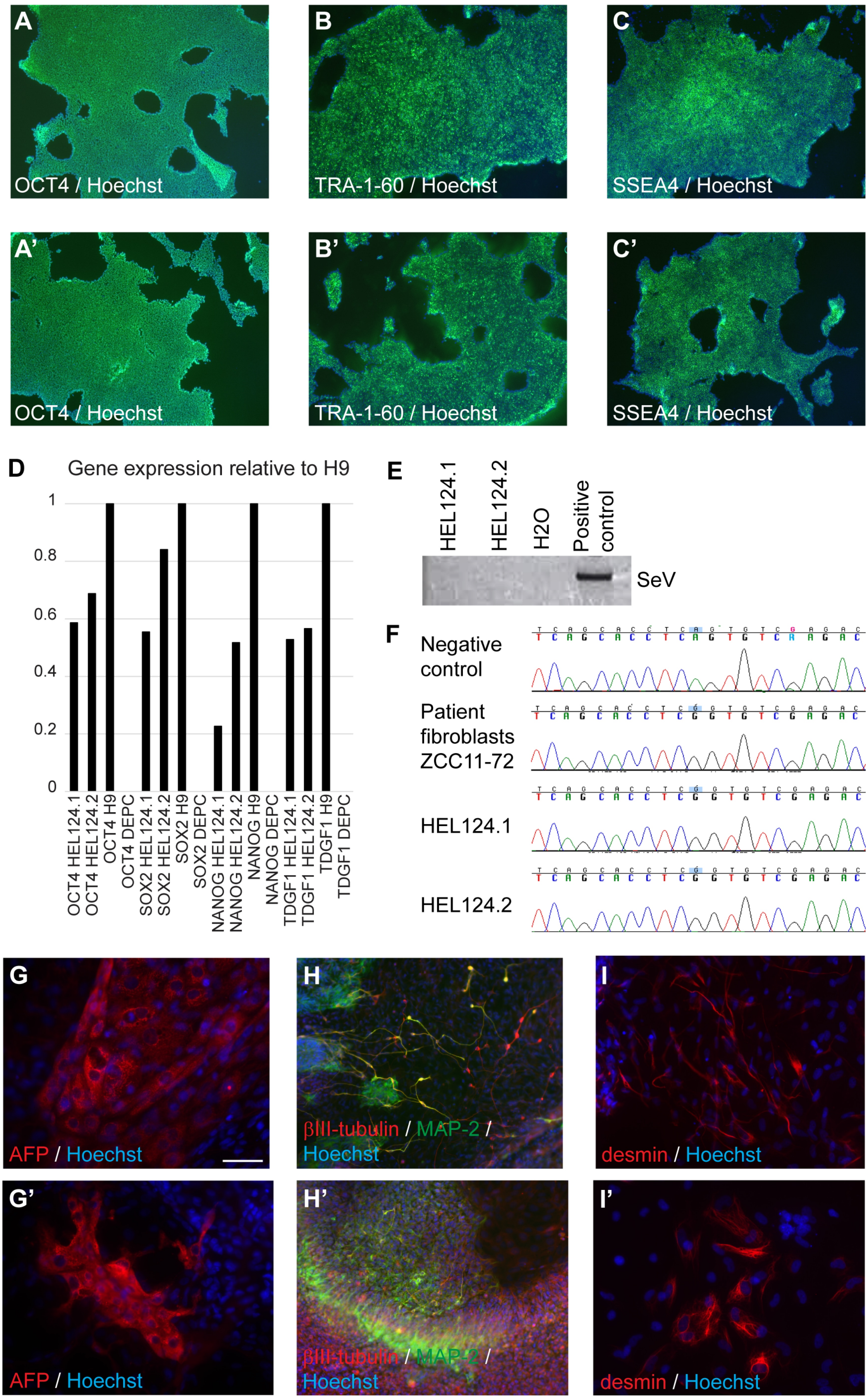
A
OCT4 / Hoechst
B
TRA-1-60 / Hoechst
C
SSEA4 / Hoechst
A'
OCT4 / Hoechst
B'
TRA-1-60 / Hoechst
C'
SSEA4 / Hoechst
D
Gene expression relative to H9
1
0.8
0.6
0.4
0.2
0
OCT4 HEL124.1
OCT4 HEL124.2
OCT4 H9
OCT4 DEPC
SOX2 HEL124.1
SOX2 HEL124.2
SOX2 H9
SOX2 DEPC
NANOG HEL124.1
NANOG HEL124.2
NANOG H9
NANOG DEPC
TDGF1 HEL124.1
TDGF1 HEL124.2
TDGF1 H9
TDGF1 DEPC
E
HEL124.1
HEL124.2
H2O
Positive control
SeV
F
Negative control
Patient fibroblasts ZCC11-72
HEL124.1
HEL124.2
G
AFP / Hoechst
H
βIII-tubulin / MAP-2 / Hoechst
I
desmin / Hoechst
G'
AFP / Hoechst
H'
βIII-tubulin / MAP-2 / Hoechst
I'
desmin / Hoechst

**Supplementary Figure S2. Characterisation of iPSC clones HEL124.1 and HEL124.2 from a newborn female GRACILE patient (ZCC11-72).** (A–C) HEL 124.1 and (A'–C') HEL124.2 iPSC colonies had typical PSC colony morphology and were positive for the nuclear/plasma membrane stem cell markers OCT4 (A and A′), TRA-1-60 (B and B′), and SSEA4 (C and C′; nuclear Hoechst staining in blue). (D) HEL 124.1 and HEL124.2 iPSCs expressed endogenous stem cell marker genes, *OCT3/4*, *SOX2*, *NANOG*, and *TDGF1* verified by RT-qPCR (hESC line H9 used as a positive control). (E) The loss of viral expression was confirmed at passage 10 by RT-PCR using primers specific for the Sendai virus (SeV). (F) The presence of the GRACILE disease-causing mutation, c.232A>G (p.S78G) was confirmed by Sanger sequencing (mutation site highlighted with blue). Pluripotency of (G–I) HEL124.1 and (G'–I') HEL124.2 was verified by staining embryoid body-derived cultures for the markers of the three germ layers, α-fetoprotein (AFP, endoderm), β-III tubulin and MAP-2 (ectoderm), and desmin (mesoderm).

A

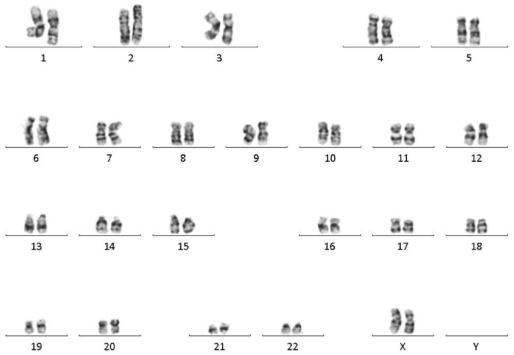


B

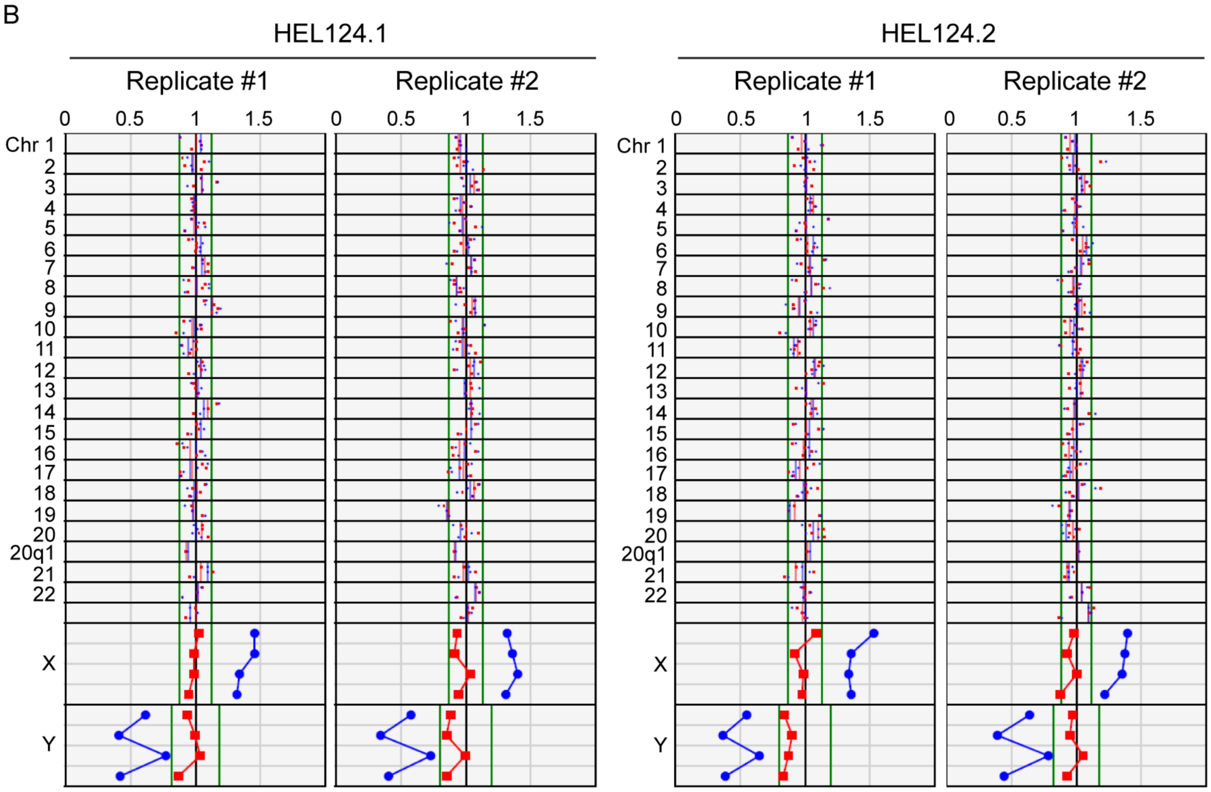


**Supplementary Figure S3. Karyotype analysis of HEL61.2 and HEL124.1 and HEL124.2 iPSC lines.** (A) Normal karyotype of HEL61.2 was determined by G-banding. (B) Chromosomal integrity of HEL124.1 and HEL124.2 was analysed by KaryoLite™ BoBs™ (data on both analysed technical replicates provided). The blue and red dots indicate the normalised chromosomal signal ratios of analysed samples against the male (blue) and female (red) reference with normal genotype. Each chromosome (Chr) was analysed with 2–3 probe sets per arm. Normal chromosomes have the signal ratios inside the reference area around value 1 while in case of chromosomal abnormalities, both signal ratios should clearly exceed the calculated threshold values in both replicates. All analysed iPSC lines were confirmed to represent females and no major chromosomal aberrations were found.

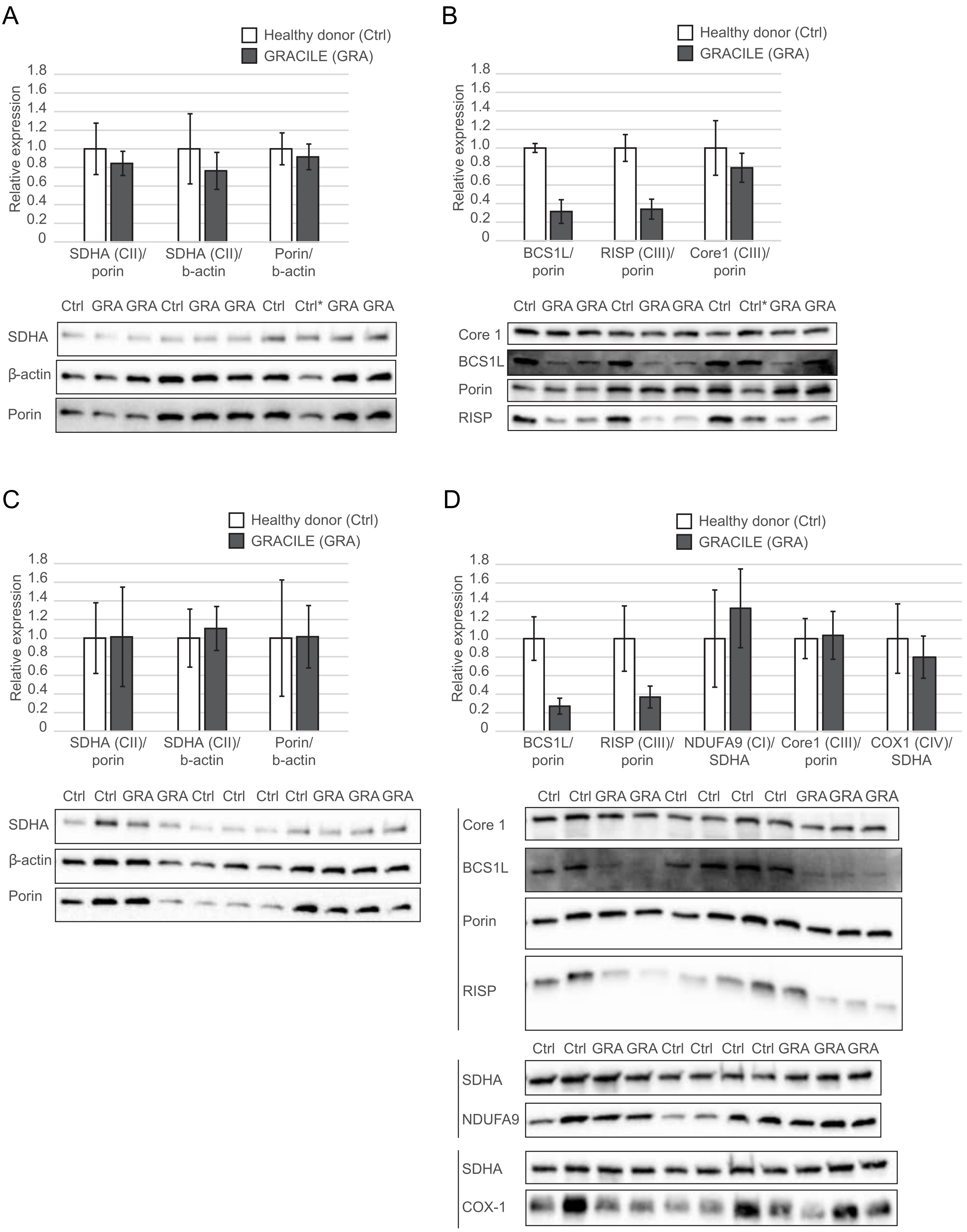
A
Healthy donor (Ctrl)
GRACILE (GRA)
Relative expression
1.8
1.6
1.4
1.2
1.0
0.8
0.6
0.4
0.2
0
SDHA (CII)/ porin
SDHA (CII)/ b-actin
Porin/ b-actin
Ctrl GRA GRA Ctrl GRA GRA Ctrl Ctrl* GRA GRA
SDHA
β-actin
Porin
B
Healthy donor (Ctrl)
GRACILE (GRA)
Relative expression
BCS1L/ porin
RISP (CIII)/ porin
Core1 (CIII)/ porin
Ctrl GRA GRA Ctrl GRA GRA Ctrl Ctrl* GRA GRA
Core 1
BCS1L
Porin
RISP
C
Healthy donor (Ctrl)
GRACILE (GRA)
Relative expression
SDHA (CII)/ porin
SDHA (CII)/ b-actin
Porin/ b-actin
Ctrl Ctrl GRA GRA Ctrl Ctrl Ctrl Ctrl GRA GRA GRA
SDHA
β-actin
Porin
D
Healthy donor (Ctrl)
GRACILE (GRA)
Relative expression
BCS1L/ porin
RISP (CIII)/ porin
NDUFA9 (CI)/ SDHA
Core1 (CIII)/ porin
COX1 (CIV)/ SDHA
Ctrl Ctrl GRA GRA Ctrl Ctrl Ctrl Ctrl GRA GRA GRA
Core 1
BCS1L
Porin
RISP
Ctrl Ctrl GRA GRA Ctrl Ctrl Ctrl Ctrl GRA GRA GRA
SDHA
NDUFA9
SDHA
COX-1

**Supplementary Figure S4. Expression of mitochondrial respiratory chain proteins in iPSC-derived tissues generated from healthy donor and GRACILE syndrome iPSCs.** (A and B) Definitive endodermal cells differentiating towards hepatocyte lineage cells. (C and D) Kidney organoids. (A and C) Since outer mitochondrial membrane protein VDAC1/porin, inner mitochondrial membrane protein SDHA, and cytoplasmic β-actin had comparable relative expression levels in controls vs affected cell lines, these loading controls were considered equivalent in the normalization of the remaining tested mitochondrial proteins as feasible in terms of molecular size and relative abundance of each target protein in iPSC-derivatives: BCS1L, RISP, and, where indicated, mitochondrial respiratory chain proteins NDUFA9 (CI), Core1 (CIII), and COX1 (CIV). Cell lines representing the same genotype (control or GRACILE) were pooled for visualization of the mean results with standard deviations. Cell lines and number of biologic replicates pooled in each assessment were as follows: healthy donor iPSCs HEL24.3 (n=3) and HEL61.2 (n=1* in A and B, n=3 in C and D), and GRACILE iPSCs HEL124.1 (n=3 in A and B) and HEL124.2 (n=3 in A and B, n=5 in C and D). Original, uncropped, unadjusted blot images are shown in Supplementary File 2.

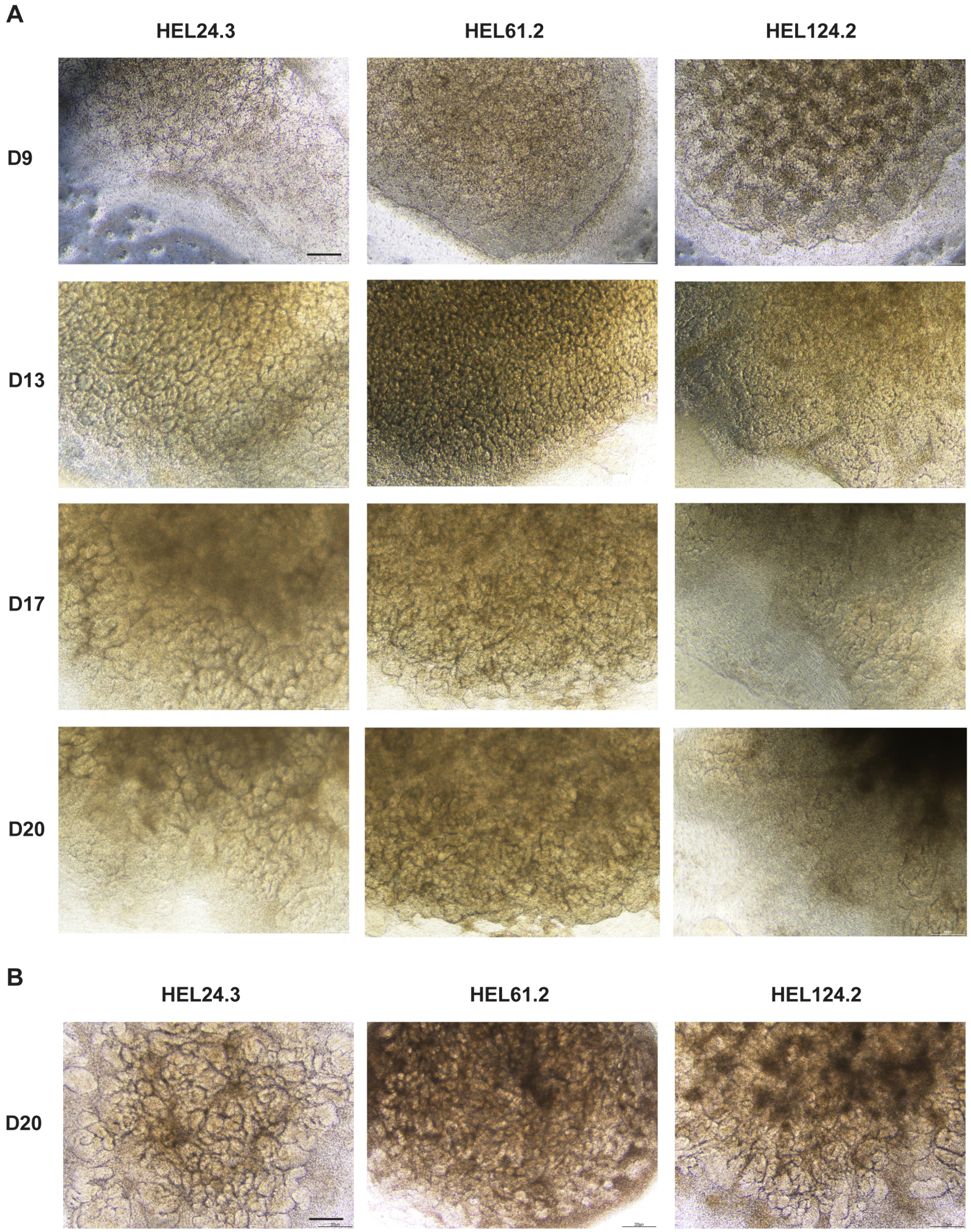


**Supplementary Figure S5. Appearance of GRACILE iPSC line HEL124.2 kidney organoids produced in the APEL/air-medium interface system.** Representative light microscopy images of kidney organoids generated from healthy donor iPSC lines (HEL24.3 and HEL61.2) and GRACILE iPSC line (HEL124.2) at indicated time points of the differentiation. (A) In the initial analysis consisting of four separate experiments, the GRACILE iPSC line generated internal structures similarly to the ones produced by the healthy donor cell lines (day 9 and 13), but subsequently, the structures started to disappear/degenerate further the differentiation was progressed resulting in an organoid with a few podocyte clusters and abnormal tubuli (day 17 and 20). (B) In four additional experiments the initial results on the phenotype were not replicated (representative images shown for day 20). Scale bar 200 µm (A–B).

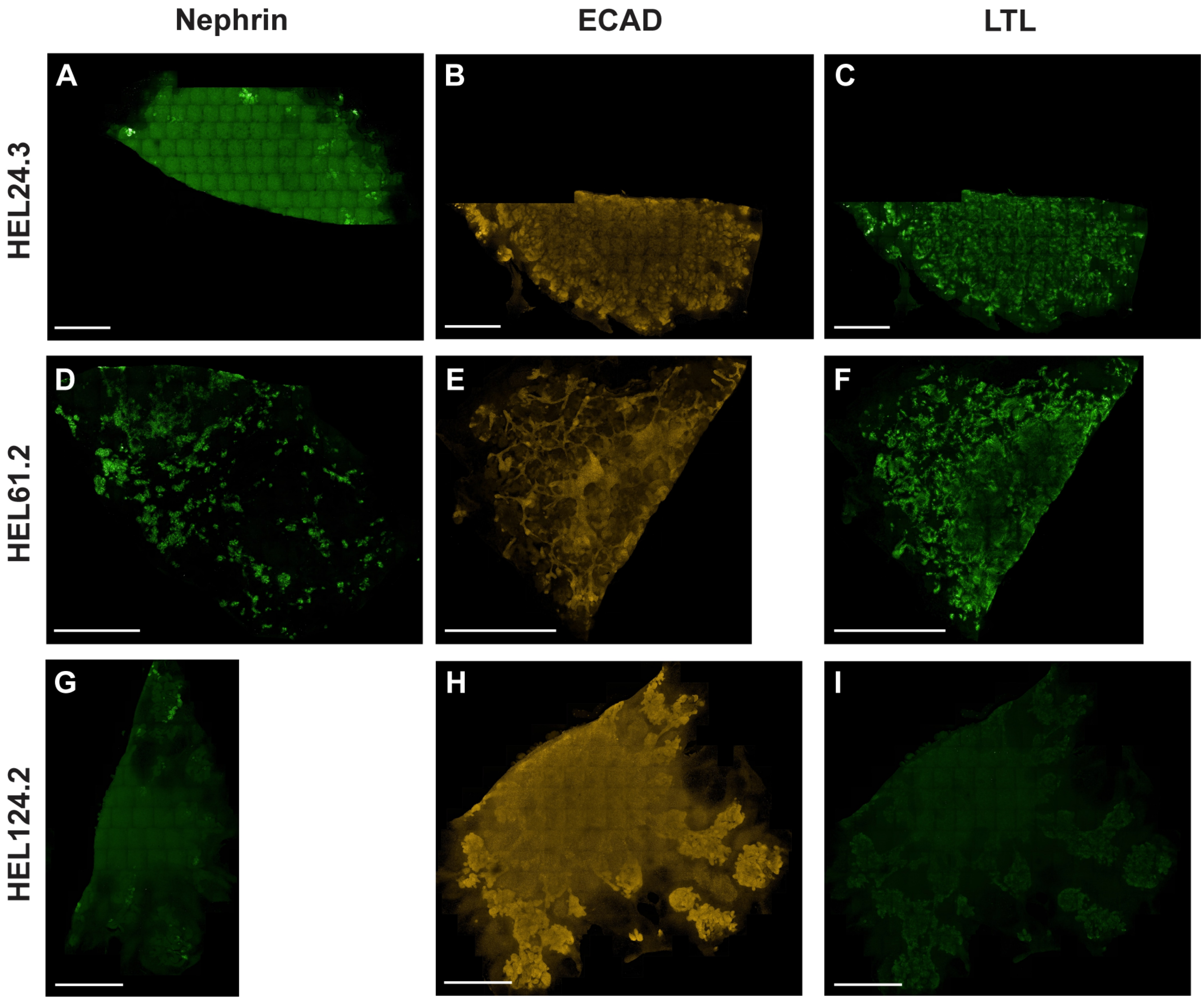


**Supplementary Figure S6. A lower magnification image of kidney organoid glomerular and tubular structures.** Kidney organoids generated with the original APEL/air-medium-interphase protocol were immunostained with a podocyte-marker (nephrin, green), a marker for maturing tubular epithelial cells, (E-cadherin, ECAD, red) and a marker for proximal tubule cells (lotus tetragonolobus lectin, LTL, green). Images acquired with the Opera Phenix spinning disk confocal microscope of organoids generated from (A-F) healthy-donor iPSC lines HEL24.3 and HEL61.2, and (G-I) the GRACILE iPSC line HEL124.2. Scale bar 1 mm.

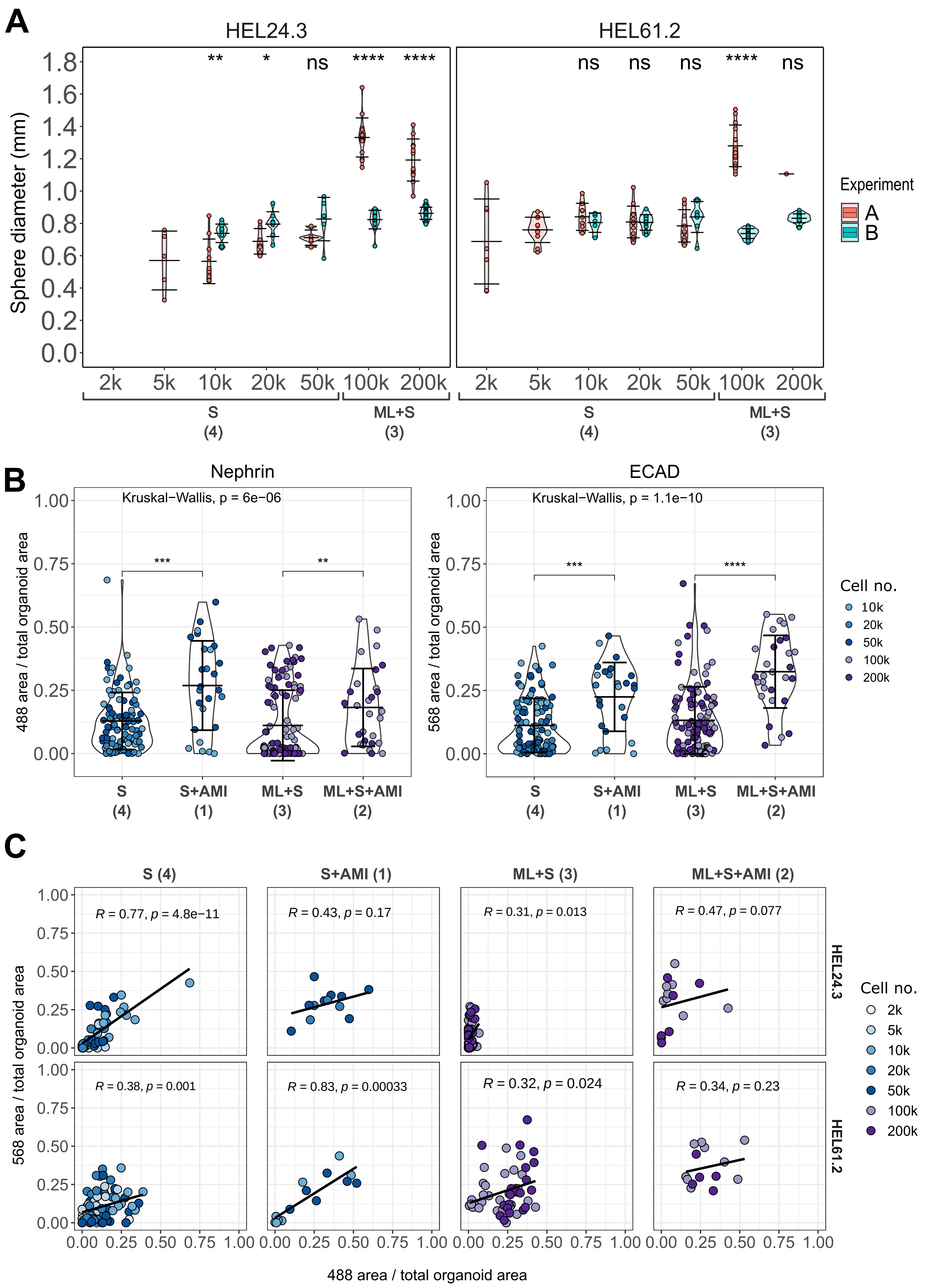
A
HEL24.3
HEL61.2
Sphere diameter (mm)
** * ns **** ****
ns ns ns **** ns
2k 5k 10k 20k 50k 100k 200k
S (4)
ML+S (3)
Experiment
A
B
B
Nephrin
ECAD
Kruskal−Wallis, p = 6e−06
Kruskal−Wallis, p = 1.1e−10
488 area / total organoid area
568 area / total organoid area
S (4)
S+AMI (1)
ML+S (3)
ML+S+AMI (2)
Cell no.
10k
20k
50k
100k
200k
C
S (4)
S+AMI (1)
ML+S (3)
ML+S+AMI (2)
R = 0.77, p = 4.8e−11
R = 0.43, p = 0.17
R = 0.31, p = 0.013
R = 0.47, p = 0.077
R = 0.38, p = 0.001
R = 0.83, p = 0.00033
R = 0.32, p = 0.024
R = 0.34, p = 0.23
HEL24.3
HEL61.2
568 area / total organoid area
488 area / total organoid area
Cell no.
2k
5k
10k
20k
50k
100k
200k

**Supplementary Figure S7. Quantification of kidney organoid size and structure formation in organoids generated with the higher throughput approaches.** Kidney organoids were generated with higher throughput approaches 1-4 (monolayer culture, ML, spheroid suspension culture, S, air-medium interface, AMI), immunostained with the podocyte marker, nephrin and tubular epithelial cell marker, E-cadherin (ECAD), and imaged with Opera Phenix high-content screening system. Organoid differentiation was conducted with two iPSC lines (HEL24.3 and HEL61.2) in two different experiments (A and B) and started with variable number of cells at the initiation of spheroids, from 2 000 (2k) to 200 000 (200k) cells. Each dot represents one organoid (mean ± s.d. indicated with black crossbars). (A) Diameter of spheroid organoids generated with higher throughput approaches 3 and 4. (B) Proportion of nephrin and ECAD-positive area per organoid is shown (pooled data from two cell lines and experiments). (A-B) Unadjusted significance levels are shown (Mann-Whitney U test) *: $p < 0.05$, **: $p < 0.01$, ***: $p < 0.001$, ****: $p < 0.0001$, ns: non-significant. (C) Correlation of nephrin and ECAD-positive area per organoid (pooled data from two experiments, Spearman's correlation coefficient; R).

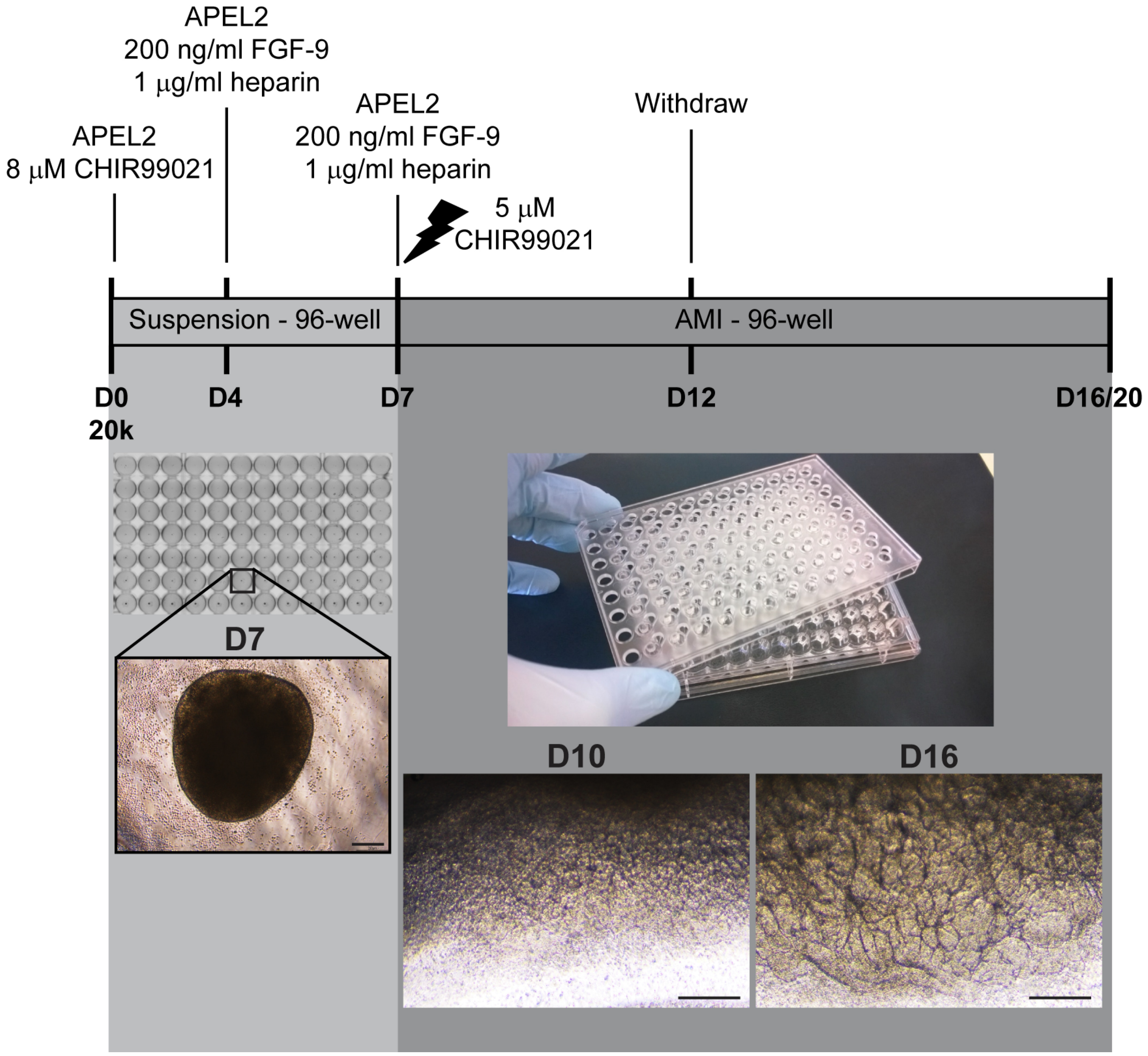


**Supplementary Figure S8. Overview of the approach for the generation of kidney organoids at air-medium interface (AMI) in APEL medium on 96-well plates.** Differentiation was started with spheroid cultures with 20 000 cells at the initiation. At day 7, spheroids were placed on Transwell membranes at AMI. Scale bars 200 µm.

### Supplementary Tables

**Supplementary Table S1. Antibodies utilized in immunocytochemistry of iPSCs and embryoid bodies.**

| Host | Type | Antigen | Company | Product number | Dilution |
|---|---|---|---|---|---|
| Rabbit | Monoclonal | Human octamer-binding transcription factor 4 (OCT4) | Cell Signaling | C30A3 | 1:500 |
| Mouse | Monoclonal | Human TRA-1-60 (podocalyxin) | Thermo Fisher | MA1-023 | 1:500 |
| Rat | Monoclonal | Mouse stage-specific embryonic antigen-3 (SSEA3) | Millipore | MAB4303 | 1:1 000 |
| Mouse | Monoclonal | Human stage-specific embryonic antigen-4 (SSEA4) | Millipore | MAB4304 | 1:1 000 |
| Rabbit | Polyclonal | Human α-fetoprotein (AFP) | Dako | A0008 | 1:400 |
| Mouse | Monoclonal | Rat TUJ-1/β-III tubulin | R&D Systems | MAB1195 | 1:500 |
| Rabbit | Polyclonal | Rat TUJ-1/β-III tubulin | Covance | PRB435P | 1:1 500 |
| Rabbit | Polyclonal | Human vimentin | Santa Cruz | sc-5565 | 1:500 |
| Rabbit | Monoclonal | Human desmin | Epitomics | 1466-1 | 1:500 |
| Donkey | Polyclonal | Goat IgG (AlexaFluor 488-conjugated) | Invitrogen | A11055 | 1:500 |
| Donkey | Polyclonal | Goat IgG (AlexaFluor 594-conjugated) | Invitrogen | 11058 | 1:500 |
| Donkey | Polyclonal | Mouse IgG (AlexaFluor 594-conjugated) | Invitrogen | A21203 | 1:500 |
| Donkey | Polyclonal | Mouse IgG (AlexaFluor 488-conjugated) | Invitrogen | A21202 | 1:500 |
| Donkey | Polyclonal | Rat IgG (AlexaFluor 488-conjugated) | Invitrogen | A-21208 | 1:500 |
| Donkey | Polyclonal | Rabbit IgG (AlexaFluor 488-conjugated) | Invitrogen | A21206 | 1:500 |
| Donkey | Polyclonal | Rabbit IgG (AlexaFluor 594-conjugated) | Invitrogen | A21207 | 1:500 |

**Supplementary Table S2. Antibodies utilized in Western blot analyses of hepatocyte lineage cells and kidney organoids.**

| Host | Type | Antigen (observed molecular size, ≈ kDa) | Company | Product number | Dilution |
|---|---|---|---|---|---|
| Mouse | Mc | Succinate dehydrogenase complex, subunit A (SDHA; 70 kDa) | Abcam | ab14715 | 1:5 000 |
| Rabbit | Pc | Human BCS1 Homolog, Ubiquinol-Cytochrome C Reductase Complex Chaperone (BCS1L; 48 kDa) | Sigma | HPA037701 | 1:1 000 |
| Mouse | Mc | Ubiquinol-cytochrome C reductase core protein 1 (Core1; 49 kDa) | Abcam | ab110252 | 1:5 000 |
| Mouse | Mc | Rieske iron-sulfur protein (RISP/UQCRFS1; 25 kDa) | Abcam | ab14746 | 1:2 500 |
| Rabbit | Mc | Human voltage-dependent anion channel 1 (VDAC1/porin; 33 kDa) | Abcam | ab154856 | 1:5 000 |
| Mouse | Mc | Cytochrome c oxidase 1 (Cox1; 33 kDa) | Abcam | ab14705 | 1:3 000 |
| Mouse | Mc | Cow NADH dehydrogenase (ubiquinone) 1 alpha subcomplex subunit 9 (NDUFA9; 36 kDa) | Abcam | ab14713 | 1:2 000 |
| Mouse | Mc | Human β-actin (45 kDa) | Cell Signaling Technology | 3700 | 1:10 000 |
| Horse | Pc | HRP-linked anti-mouse IgG | Cell Signaling Technology | 7076S | 1:2 000–1:5 000 |
| Goat | Pc | HRP-linked anti-rabbit IgG | Cell Signaling Technology | 7074S | 1:2 000–1:5 000 |

Abbreviations: HRP, horseradish peroxidase; Mc, monoclonal; Pc, polyclonal

**Supplementary Table S3. Kidney organoids generated with higher throughput approaches and imaged with Opera Phenix (number of replicates per condition).**

| Experiment A/B | Cell line | Marker | Approach | Cell number (k, 1000 cells) at the initiation of spheroids | n |
|---|---|---|---|---|---|
| A | HEL24.3 | nephrin/ECAD | ML+S+AMI | 100k | 3 |
| A | HEL24.3 | nephrin/ECAD | ML+S+AMI | 200k | 3 |
| A | HEL24.3 | nephrin/ECAD | S+AMI | 20k | 2 |
| A | HEL24.3 | nephrin/ECAD | S+AMI | 50k | 3 |
| A | HEL24.3 | nephrin/ECAD | ML+S | 100k | 15 |
| A | HEL24.3 | nephrin/ECAD | ML+S | 200k | 12 |
| A | HEL24.3 | nephrin/ECAD | S | 5k | 5 |
| A | HEL24.3 | nephrin/ECAD | S | 10k | 10 |
| A | HEL24.3 | nephrin/ECAD | S | 20k | 7 |
| A | HEL24.3 | nephrin/ECAD | S | 50k | 5 |
| A | HEL61.2 | nephrin/ECAD | ML+S+AMI | 100k | 3 |
| A | HEL61.2 | nephrin/ECAD | ML+S+AMI | 200k | 3 |
| A | HEL61.2 | nephrin/ECAD | S+AMI | 10k | 3 |
| A | HEL61.2 | nephrin/ECAD | S+AMI | 50k | 3 |
| A | HEL61.2 | nephrin/ECAD | ML+S | 100k | 14 |
| A | HEL61.2 | nephrin/ECAD | ML+S | 200k | 1 |
| A | HEL61.2 | nephrin/ECAD | S | 2k | 7 |
| A | HEL61.2 | nephrin/ECAD | S | 5k | 11 |
| A | HEL61.2 | nephrin/ECAD | S | 10k | 9 |
| A | HEL61.2 | nephrin/ECAD | S | 20k | 12 |
| A | HEL61.2 | nephrin/ECAD | S | 50k | 9 |
| B | HEL24.3 | nephrin/ECAD | ML+S+AMI | 100k | 6 |
| B | HEL24.3 | nephrin/ECAD | ML+S+AMI | 200k | 3 |
| B | HEL24.3 | nephrin/ECAD | S+AMI | 20k | 2 |
| B | HEL24.3 | nephrin/ECAD | S+AMI | 50k | 5 |
| B | HEL24.3 | nephrin/ECAD | ML+S | 100k | 15 |
| B | HEL24.3 | nephrin/ECAD | ML+S | 200k | 21 |
| B | HEL24.3 | nephrin/ECAD | S | 10k | 9 |
| B | HEL24.3 | nephrin/ECAD | S | 20k | 8 |
| B | HEL24.3 | nephrin/ECAD | S | 50k | 6 |
| B | HEL61.2 | nephrin/ECAD | ML+S+AMI | 100k | 6 |
| B | HEL61.2 | nephrin/ECAD | ML+S+AMI | 200k | 2 |
| B | HEL61.2 | nephrin/ECAD | S+AMI | 10k | 5 |
| B | HEL61.2 | nephrin/ECAD | S+AMI | 50k | 3 |
| B | HEL61.2 | nephrin/ECAD | ML+S | 100k | 16 |
| B | HEL61.2 | nephrin/ECAD | ML+S | 200k | 20 |
| B | HEL61.2 | nephrin/ECAD | S | 10k | 5 |
| B | HEL61.2 | nephrin/ECAD | S | 20k | 12 |
| B | HEL61.2 | nephrin/ECAD | S | 50k | 8 |
